# Species-scale genomic analysis of *S. aureus* genes influencing phage host range and their relationships to virulence and antibiotic resistance genes

**DOI:** 10.1101/2021.08.23.457453

**Authors:** Abraham G. Moller, Robert A. Petit, Timothy D. Read

**Affiliations:** Division of Infectious Diseases, Department of Medicine, School of Medicine, Emory University, Atlanta, GA, USA; Department of Human Genetics, School of Medicine, Emory University, Atlanta, GA, USA

## Abstract

Phage therapy has been proposed as a possible alternative treatment for infections caused by the ubiquitous bacterial pathogen *Staphylococcus aureus.* However, successful phage therapy requires knowing both host and phage genetic factors influencing host range for rational cocktail formulation. To further our understanding of host range, we searched 40,000+ public *S. aureus* genome sequences for previously identified phage resistance genes. We found that phage adsorption targets and genes that block phage assembly were significantly more conserved than genes targeting phage biosynthesis. Core phage resistance genes had similar nucleotide diversity, ratio of non-synonymous to synonymous substitutions, and functionality (measured by delta-bitscore) to other core genes in a set of 380 non-redundant *S. aureus* genomes (each from a different MLST sequence type). Non-core phage resistance genes were significantly less consistent with the core genome phylogeny than all non-core genes in this set. Only superinfection immunity genes correlated with empirically determined temperate phage resistance, accessory genome content, and numbers of accessory antibiotic resistance or virulence genes encoded per strain. Taken together, these results suggested that, while phage adsorption genes are heavily conserved in the *S. aureus* species, they are not undergoing positive selection, arms race dynamics. They also suggested genes classified as involved in assembly are least phylogenetically constrained and superinfection immunity genes best predict both empirical phage resistance and levels of phage-mediated HGT.

**Importance:** *Staphylococcus aureus* is a widespread, hospital- and community-acquired pathogen that is commonly antibiotic resistant. It causes diverse diseases affecting both the skin and internal organs. Its ubiquity, antibiotic resistance, and disease burden make new therapies urgent, such as phage therapy, in which viruses specific to infecting bacteria clear infection. *S. aureus* phage host range not only determines whether phage therapy will be successful by killing bacteria but also horizontal gene transfer through transduction of host genetic material by phages. In this work, we comprehensively reviewed existing literature to build a list of *S. aureus* phage resistance genes and searched our database of almost 43,000 *S. aureus* genomes for these genes to understand their patterns of evolution, finding that prophages’ superinfection immunity correlates best with phage resistance and HGT. These findings improved our understanding of the relationship between known phage resistance genes and phage host range in the species.

## Introduction

New treatments are needed for *Staphylococcus aureus* infections because of its high prevalence, increasing antibiotic resistance, diverse pathologies, and a lack of available vaccines. Phage therapy - clearing infecting bacteria with bacteriophages has been proposed as a possible alternative treatment. Potential phage therapy advantages over antibiotics include reduced toxicity because of the specificity of the virus for its host and the high diversity of natural phages available to be isolated for treatment, lessening chances of development of complete resistance (1, 2). However, no natural phage is known to kill all *S. aureus* strains, making phage cocktails (combinations of phages that have non-overlapping host ranges) necessary for successful treatment. Comprehensive knowledge of host and phage genetic factors influencing host range is needed for rational cocktail formulation.

Known *S. aureus* phages, which all belong to the order *Caudovirales* (tailed phages), are divided into three morphological classes - the long, noncontractile-tailed *Siphoviridae*, the long, contractile-tailed *Myoviridae*, and the short, noncontractile-tailed *Podoviridae* (3). The *Siphoviridae* are both temperate and virulent, while the *Myo*- and *Podoviridae* are virulent (3). The *Siphoviridae* bind either ɑ-O-GlcNAc or β-O-GlcNAc attached at the 4 positions of wall teichoic acid (WTA) ribitol phosphate monomers, the *Podoviridae* bind only β-O-GlcNAc-decorated WTA, and the *Myoviridae* bind either the WTA ribitol-phosphate backbone or β-O-GlcNAc-decorated WTA (4–6).

Bacteriophage life-cycles have been proposed to consist of five phages: attachment, uptake, biosynthesis, assembly, and lysis (7, 8). From extensive literature review, we found that reported *Staphylococcus* phage resistance mechanisms only acted at the adsorption, biosynthesis, and assembly infection stages of the phage life cycle (4). Adsorption resistance mechanisms include phage receptor alteration, removal, or occlusion by large surface proteins or polysaccharides (capsule) (5, 9–14). Biosynthesis resistance mechanisms involve halting infection through metabolic arrest (abortive infection) or phage DNA degradation by adaptive (CRISPR) or innate (restriction-modification) immunity (15–19). Assembly resistance occurs by assembly interference, in which *Staphylococcus aureus* pathogenicity islands (SaPIs), chromosomal phage-like elements, divert away the assembly of helper infecting viruses (*Siphoviridae*) toward their own, enabling them to replicate at the cost of helper viruses (20–25). In this work, we searched 40,000+ annotated *S. aureus* genomes for genes known to influence phage resistance and thus potentially influence host range. We wanted to investigate the relationships between predicted phage resistance, empirically determined phage resistance, and other genes frequently transferred between *S. aureus by* horizontal gene transfer (HGT). We developed scoring metrics for resistance determinant diversity, abundance (frequency amongst strains), functionality, and overlap, as well as overall predicted phage resistance per strain, and accessory gene content as a measure of horizontal gene transfer. We then evaluated the correlation between genome-predicted phage resistance and either empirically determined resistance levels, levels of horizontal gene transfer, or networks of gene transfer amongst strains. We anticipate the conclusions of this work will both improve phage resistance prediction, thus improving phage therapy potential, and understanding the evolution of the *S. aureus* species by elucidation of genetic determinants affecting HGT.

## Results

### Distribution of genes influencing phage resistance in S. aureus genomes

We curated a list of 331 genes located on bacterial chromosomes and plasmids (Supplemental Table S1) implicated in previous studies to influence phage resistance. Genes were included based on three criteria: 1) they were directly shown to influence phage resistance in a *S. aureus* strain in laboratory-based studies, 2) *S. aureus* phage resistant mutants had mutations in the gene, 3) they were shown to be associated with phage resistance in other bacterial species. We found that the overwhelming majority of genes “positively” influenced resistance to phages (306/331), i.e., they had the potential to increase the bacterial cell survival. A smaller number (25/331) were of potentially negative effect, i.e., bacteria may become more sensitive to phages. Each gene was classified as interfering with either the attachment (87/331), biosynthesis (235/331) or assembly (8/331) portion of the phage life cycle.

We first determined whether each gene was core or non-core within the *S. aureus* pangenome based on a search of the Staphopia database containing nearly 43,000 *S. aureus* genomes (26). We found that the most conserved (core) genes in the database were mainly adsorption genes (Figure 1A; 51/63 core genes present in over 80% of Staphopia genomes). The least conserved genes (less than 100 strain matches) were either biosynthesis or adsorption genes (Figure 1B). Both adsorption resistance and assembly resistance genes were significantly (p<0.05) more conserved than biosynthesis resistance genes (hereafter referred to by resistance category - e.g., adsorption) but adsorption genes did not significantly differ in conservation from assembly genes (Figure 1C) based on non-parametric Wilcoxon tests. Conversely, no assembly resistance genes were core genes (Figure 1A).

**Figure 1:**
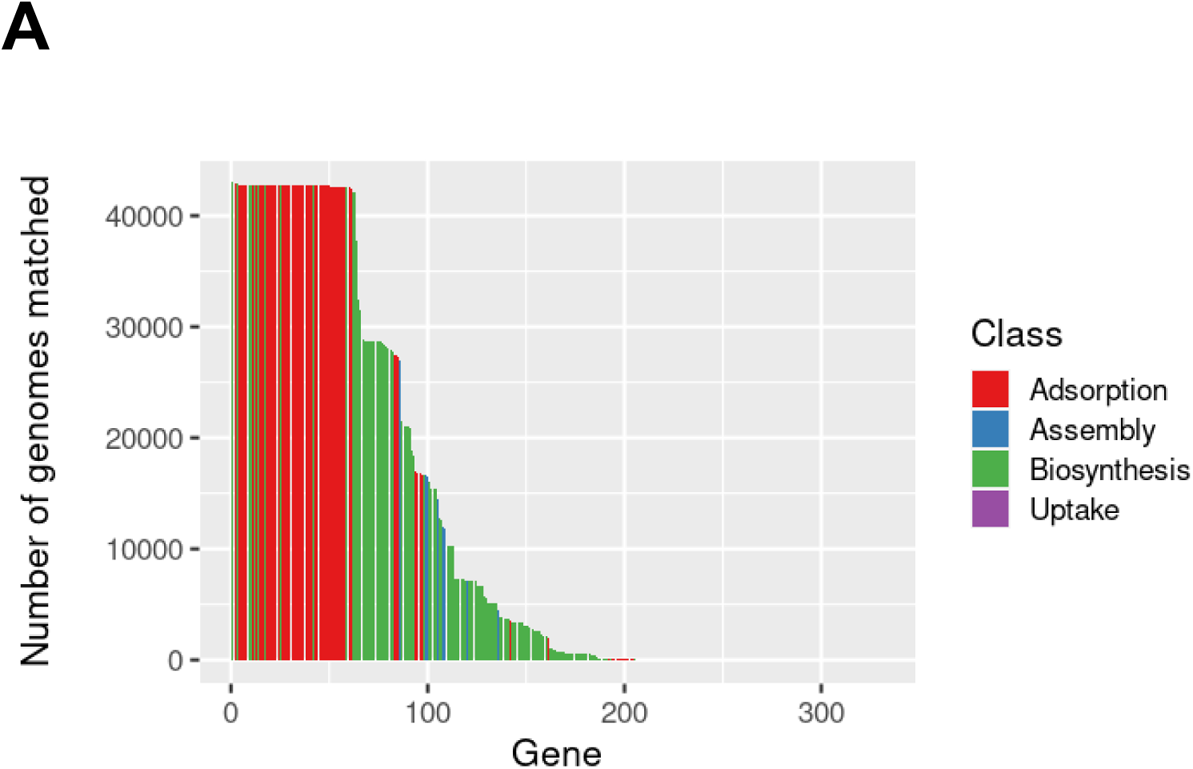
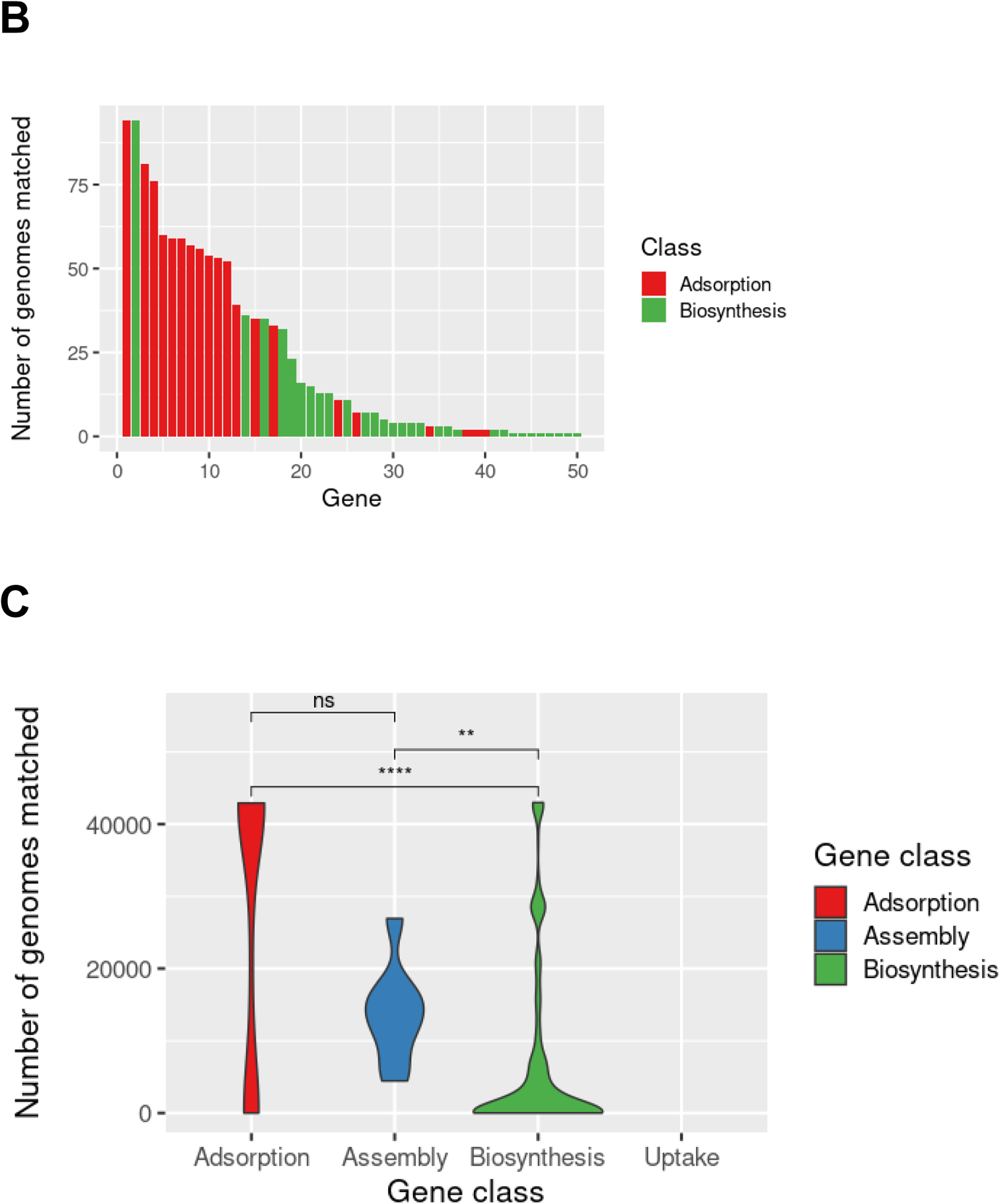
Conservation of examined phage resistance genes in the species based on a search of 40,000+ *S. aureus* genomes. We used BLAST to search for our set of 331 curated phage resistance genes in the Staphopia database. A) Conservation of each gene (y-axis) ranked from most to least conserved on the x-axis. Genes are colored by class - adsorption in red, assembly in blue, and biosynthesis in green. B) An inset of A showing genes with 100 or fewer strain matches is included in the upper right hand corner. C) Distributions of conservation for each considered category visualized as violin plots. Groups were tested for statistically significant differences with the non-parametric Wilcoxon test (ns, not significant; *, *P* = 0.01 to 0.05; **, *P* = 0.001 to 0.01; ***, *P* = 0.0001 to 0.001; ****, *P* = 0 to 0.0001).

The process of collating putative phage resistance genes and matching them against a large number of genomes revealed the presence of a number systems that were novel in *S. aureus*. Rare systems newly discovered in the species, listed with number of strains matching and taxon of original gene, included *abiR,* encoding an abortive infection system that inhibits phage DNA replication (27) (13 strains); *ietS,* encoding the serine protease toxin of the *IetAS* toxin-antitoxin system (28), (94 strains), and *ptuA,* encoding the ATPase component of the Septu phage defense system (29) (504 strains). Systems found in many more strains but newly reported here included Ec67/retron-TOPRIM (15372 strains), retron-TIR (15992 strains), and *avs1a*/MBL + protease-STAND (42123 strains) (Supplemental Table S1). Retrons are RNA-DNA chimeric molecules synthesized by reverse transcriptases that induce cell death upon sensing infection (30, 31), while Avs1a is an AVAST (antiviral ATPases/NTPases of the STAND superfamily) protein (32). Although these genes are common in the species, their function was only recently discovered (30, 32). We further examined the genomic contexts of non-core genes across 535 *S. aureus* complete genomes. We found these seven non-core genes (Supplemental Figure S1A-G) to be present near genes encoded by transposons as well as the chromosomally integrated mobile genetic element SCC*mec* (Supplemental Figure S1C) and restriction-modification genes (Supplemental Figure S1A, C, and D). This suggests these genes are readily transferred horizontally and associated with regions of genomic plasticity or previously known phage resistance determinants.

### Diversity, functionality, and strength of selection of core phage resistance genes was similar to core genes in general

We next assessed the diversity and functionality of core phage resistance genes in the 380 genome Staphopia non-redundant diversity (NRD) set (26). The NRD set includes one randomly-selected representative genome for every known *S. aureus* sequence type. We asked whether core genes differ in levels of diversity (measured as allele count or translated nucleotide diversity), functionality (measured as delta-bit score - difference between reference and query gene matches to a profile Hidden Markov model, where mutations at conserved amino acids have stronger effects than those at nonconserved amino acids) (33), or selection (measured as nonsynonymous to synonymous change ratio - dN/dS) from respective phage resistance genes. If phages and hosts existed in an arms race scenario, we would expect core phage resistance genes to be undergoing diversifying selection (higher average dN/dS than core genes). We also would expect decreased functionality (increased delta-bit score) in cases where inactivating genes would lead to resistance. We instead found similar diversity, functionality, and dN/dS between core genes and corresponding phage resistance gene subsets (p>0.05 with non-parametric Wilcoxon tests) (Figure 2).

**Figure 2:**
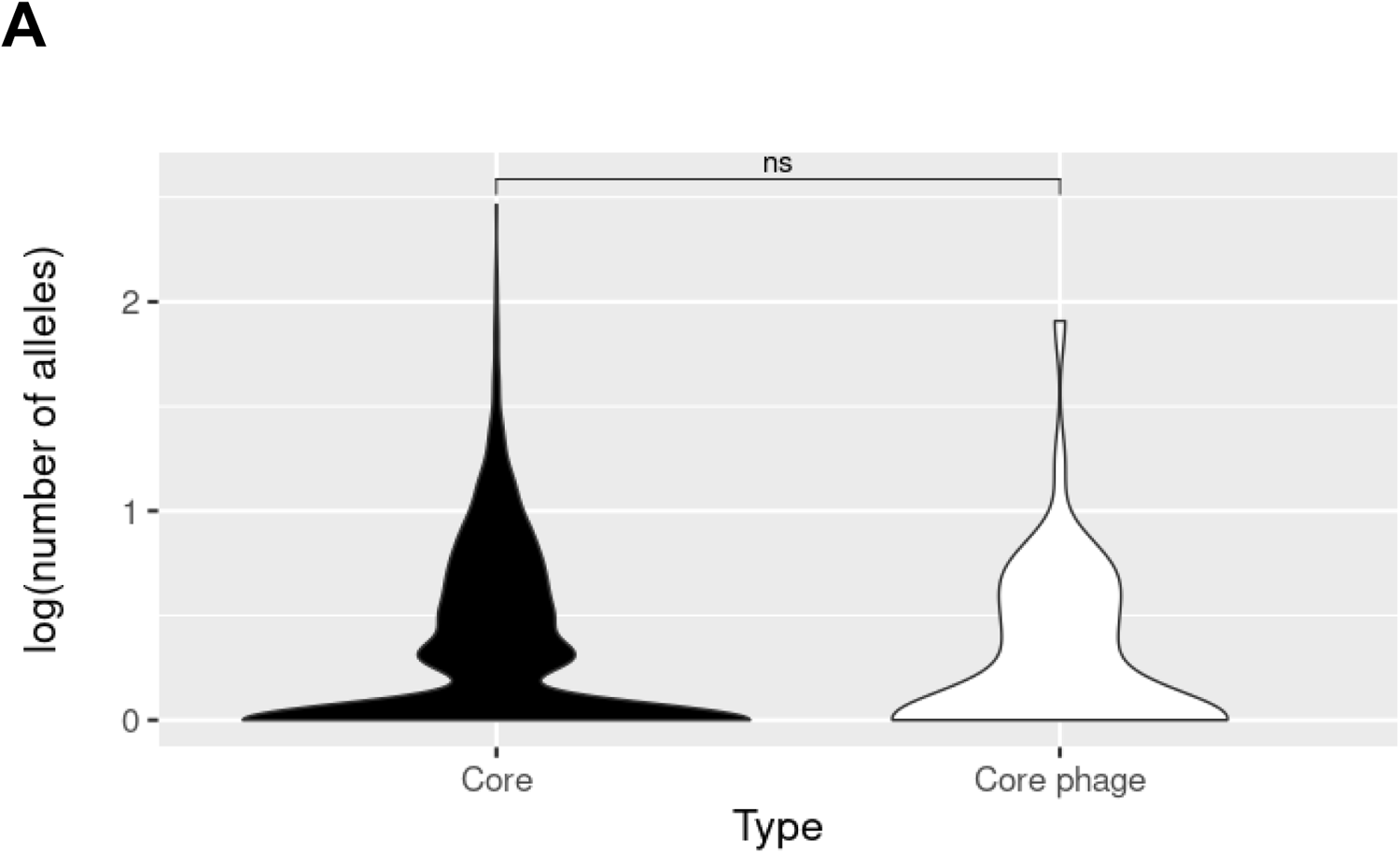
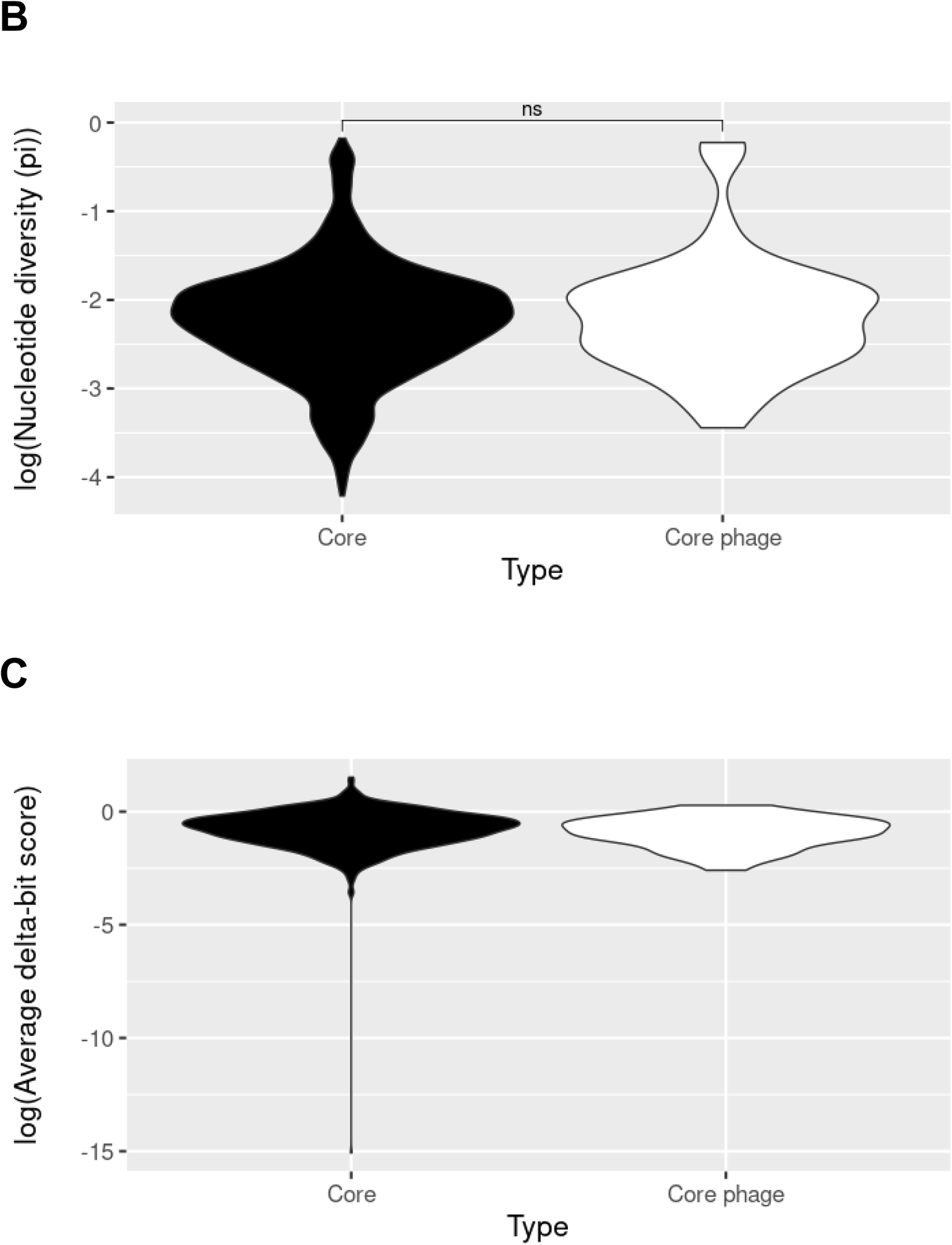
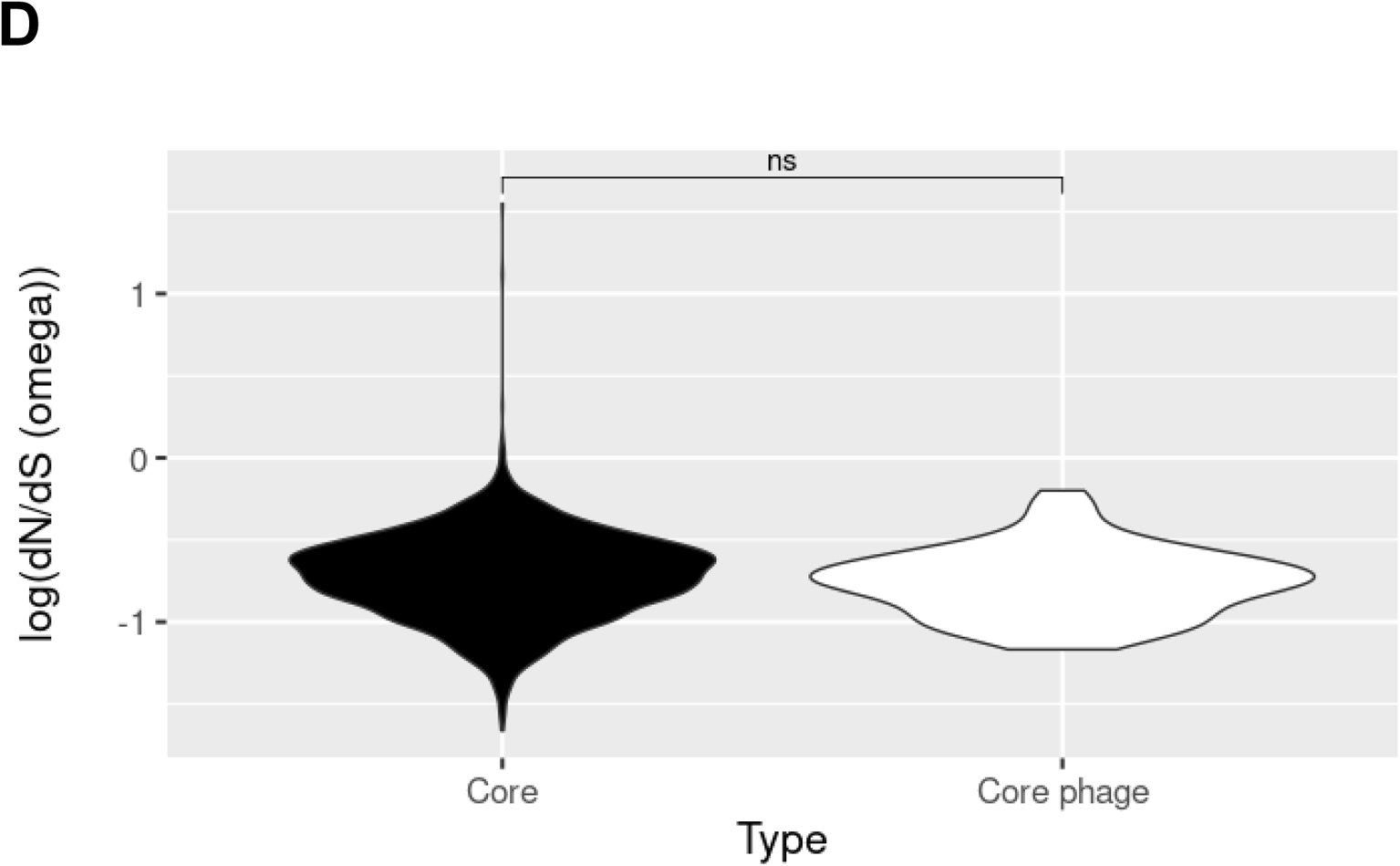
Core phage resistance genes do not differ from core genes overall in terms of diversity, functionality, or selection. Core genes were those found in 80% or more of the NRD set. Diversity (A, B) was measured as the number of alleles or translated nucleotide diversity (π). Functionality (C) was measured by delta-bit score, or calculated difference from reference profile Hidden Markov models (HMMs), which score changes from conserved amino acids higher than those from non-conserved amino acids. Selection (D) was measured through the dN/dS metric, which was calculated from gene phylogenetic trees using single-likelihood ancestor counting (SLAC). Group distributions were visualized as violin plots and differences were tested for significance using the non-parametric Wilcoxon test (ns, not significant; *, *P* = 0.01 to 0.05; **, *P* = 0.001 to 0.01; ***, *P* = 0.0001 to 0.001; ****, *P* = 0 to 0.0001). Delta-bit scores did provide enough data points for a Wilcoxon or t-test to be conducted.

### Non-core phage resistance genes have greater phylogenetic signal than other non-core genes

Phylogenetic signal is the tendency of a trait to follow the pattern of an organismal phylogenetic tree (34). The phylogenetic signal may reveal features about patterns of acquisition and uptake of a non-core gene. We asked whether non-core phage resistance genes were unusual in their phylogenetic signal compared to other non-core genes by calculating consistency indices, which measure how much gene presence/absence diverges from that expected by the phylogeny (in this case, a core genome phylogeny of the NRD set). Genes on mobile genetic elements that were frequently gained through horizontal gene transfer and lost frequently through deletion/replacement would be expected to have low consistency indices. Homoplasy - the independent loss or gain of a trait in separate lineages during evolution - is inversely proportional to consistency index and phylogenetic signal. To examine these relationships, we plotted the number of NRD strains encoding the gene of interest against the number of changes necessary, both for actual cases and the average of 999 gene presence-absence permutations per gene (Figure 3A). As expected, the permuted gene presence-absence data generated a parabola with a peak at intermediate-frequency genes. All observed gene change counts were below those expected by the parabola, indicating all genes had a level of phylogenetic signal. We found assembly genes had the highest changes amongst any group considered (Figure 3), though non-core adsorption and biosynthesis also were significantly less consistent with the phylogeny than non-core genes in general (Figure 3B). We also noted that gene frequency distribution could have an effect on the results (Supplemental Figure S2). Non-core genes in general were far less common (p<0.05) than any other category, and we showed low frequency genes under permutation tests had fewer changes and thus higher phylogenetic signal (Figure 3A) than intermediate frequency genes solely based on frequency in the genome set.

**Figure 3:**
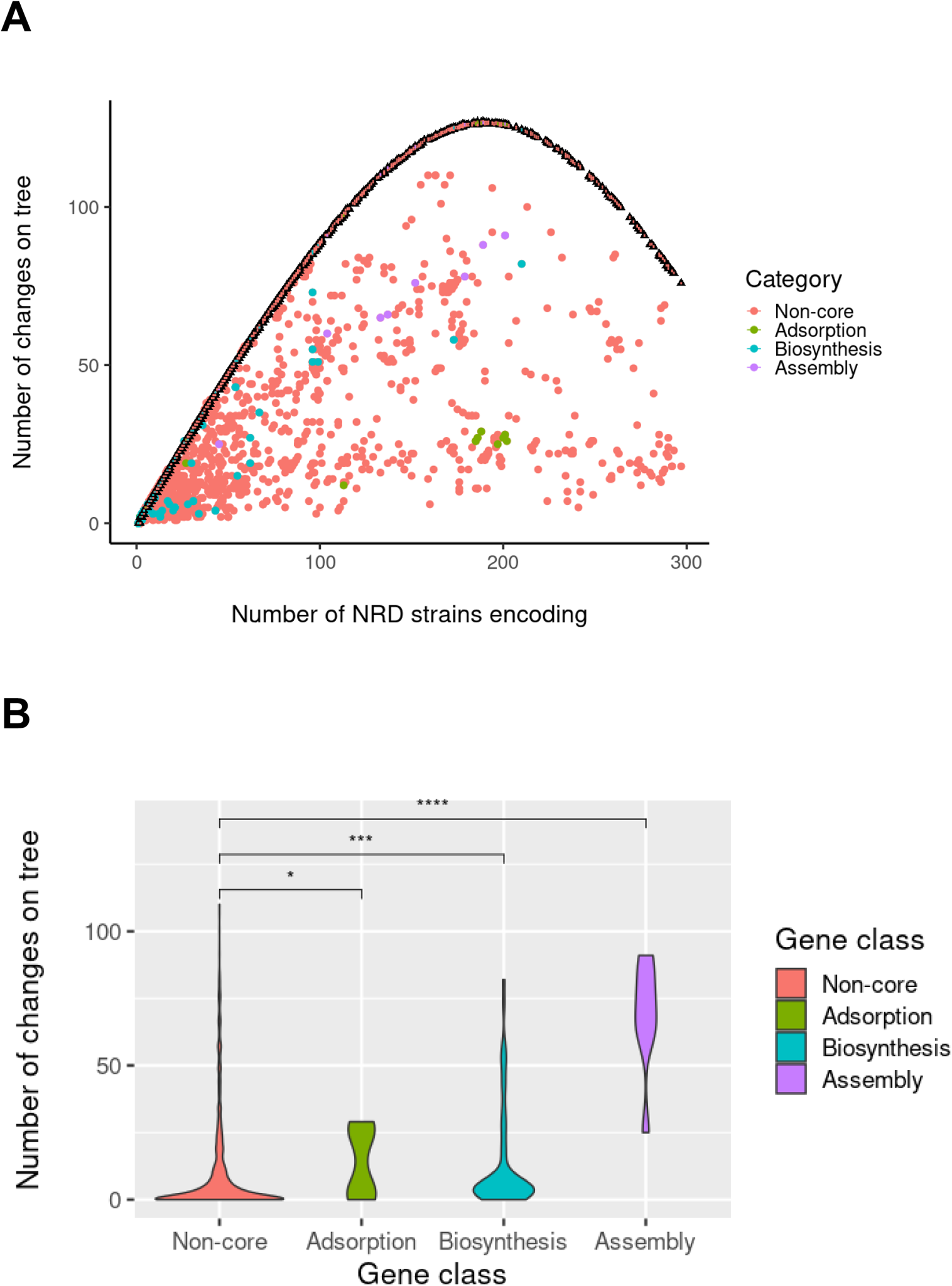
All non-core phage resistance genes have phylogenetic signal, but assembly genes have the least phylogenetic signal amongst all. We calculated phylogenetic signal as consistency index (CI) between non-core (genes present in 80% or less than the NRD set) A) The relationship between number of changes necessary to make gene presence/absence patterns consistent with the phylogeny (y-axis) and the number of strains encoding each gene (x-axis). Average change numbers after 999 permutations of gene presence/absence on the tree are shown as triangles with thick black borders and error bars (1 standard error above and below the mean), while change numbers for actual gene presence/absence are shown as circles. Non-core genes are colored in salmon red, while adsorption, biosynthesis, and assembly genes are colored in olive green, turquoise, and purple, respectively. B) Distributions of gene presence/absence changes necessary to make them consistent with the phylogeny for each gene category (non-core, adsorption, biosynthesis, and assembly) visualized as violin plots. Group differences were tested for significance using the non-parametric Wilcoxon test (ns, not significant; *, *P* = 0.01 to 0.05; **, *P* = 0.001 to 0.01; ***, *P* = 0.0001 to 0.001; ****, *P* = 0 to 0.0001).

Taken together, these results indicate all non-core phage resistance genes had lesser phylogenetic signal than other non-core genes with assembly genes having lowest values. Gene in the assembly category are frequently found on mobile SaPIs, which are frequently exchanged by HGT and lost through deletion. Unlike assembly genes, some non-core adsorption and biosynthesis genes approach complete consistency with the tree, but the proportion is lower than other non-core genes. However, we can’t discount the possibility that horizontal transfer from other *S. aureus* clades unrepresented in our database or other species could lead to rare genes completely consistent with the phylogeny.

We further examined the relationship between phylogeny and phage resistance genes by assessing the differences between clonal complexes (CC) in their numbers of non-core phage genes. For all gene categories examined, there was statistically significant variation (p<0.05) as determined by an analysis of variance (ANOVA) test (Figure 4). This suggests clonal complex is associated with both accessory genome and numbers of phage resistance genes.

**Figure 4:**
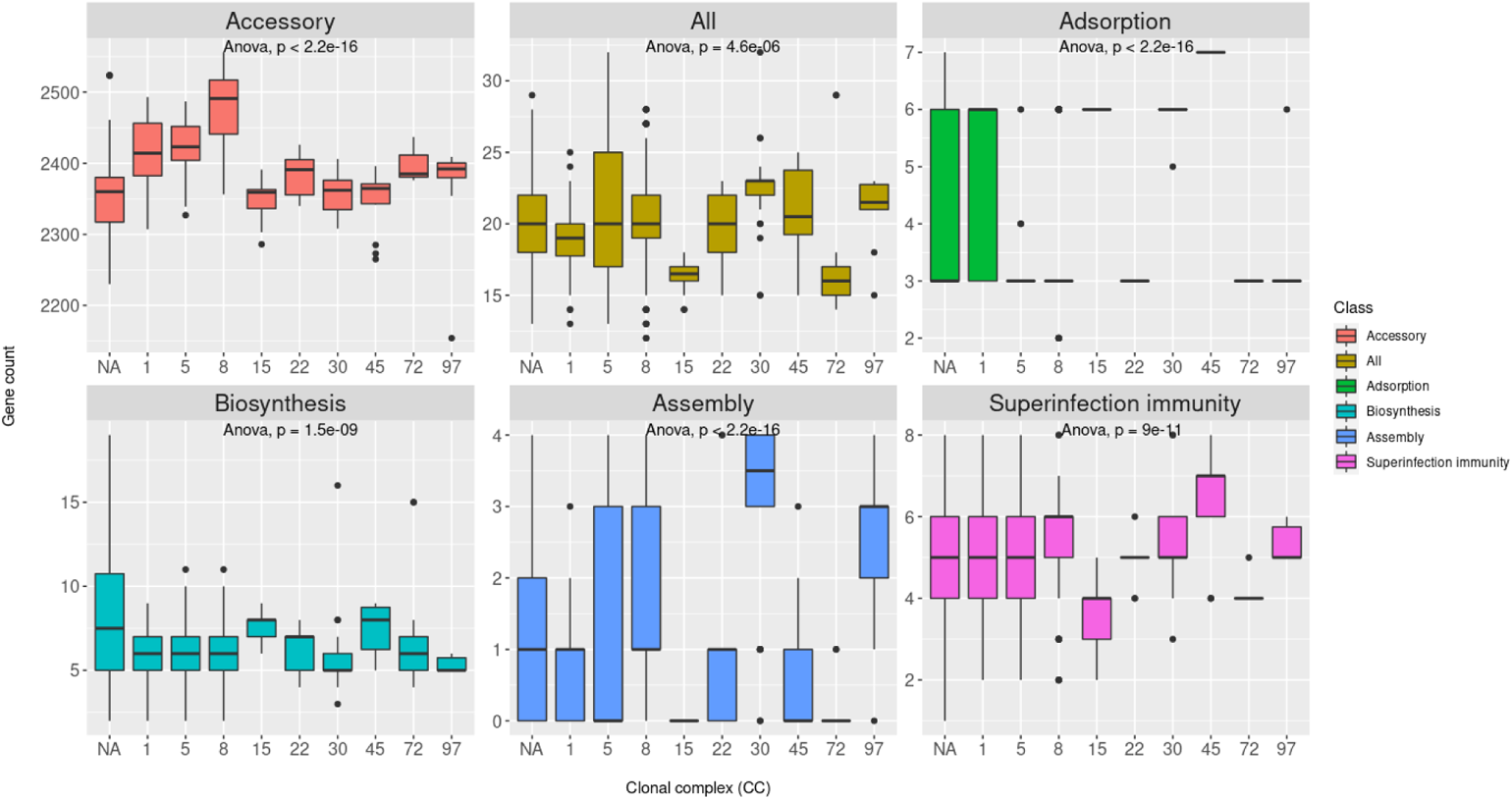
Relationship between clonal complex (CC) and accessory genome and non-core phage resistance gene content. We used BLAST to search for our set of non-core phage resistance genes and accessory genes in the set of 535 complete *S. aureus* genomes in Staphopia. We then visualized the distributions of genes for each CC as boxplots. Strains without a defined CC are listed as NA. Analysis of variance (ANOVA) assessed significant overall differences, with significance values posted on each facet.

### Non-core phage resistance genes are most often co-encoded with those of the same class

We next examined the modularity of phage resistance genes by examining “genomic overlap” in each category. We calculated genomic overlap as the average number of genes in a particular category encoded by strains containing the query gene. We hypothesized that phage resistance genes with shared functions are encoded together on the genome. We expect that such genes with shared functions are co-encoded more often than any gene in the genome at random. We found that all phage resistance genes (non-core, all phage resistance, biosynthesis) or biosynthesis genes (non-core, all phage resistance, adsorption, biosynthesis, assembly) were most often significantly different (non-parametric Wilcoxon test, p<0.05) from non-core genes (Figure 5) in subject genomic overlap distribution (subject listed here in parentheses). In each phage resistance gene category, when query and subject were the same class, query genomic overlap was most significantly different (p < 0.0001) with non-core query overlap, and increased on average in all cases (Figure 5). This supported our hypothesis that genes with shared functions were encoded together, as adsorption, biosynthesis, and assembly genes were encoded together significantly more often than similar non-core genes at random.

**Figure 5:**
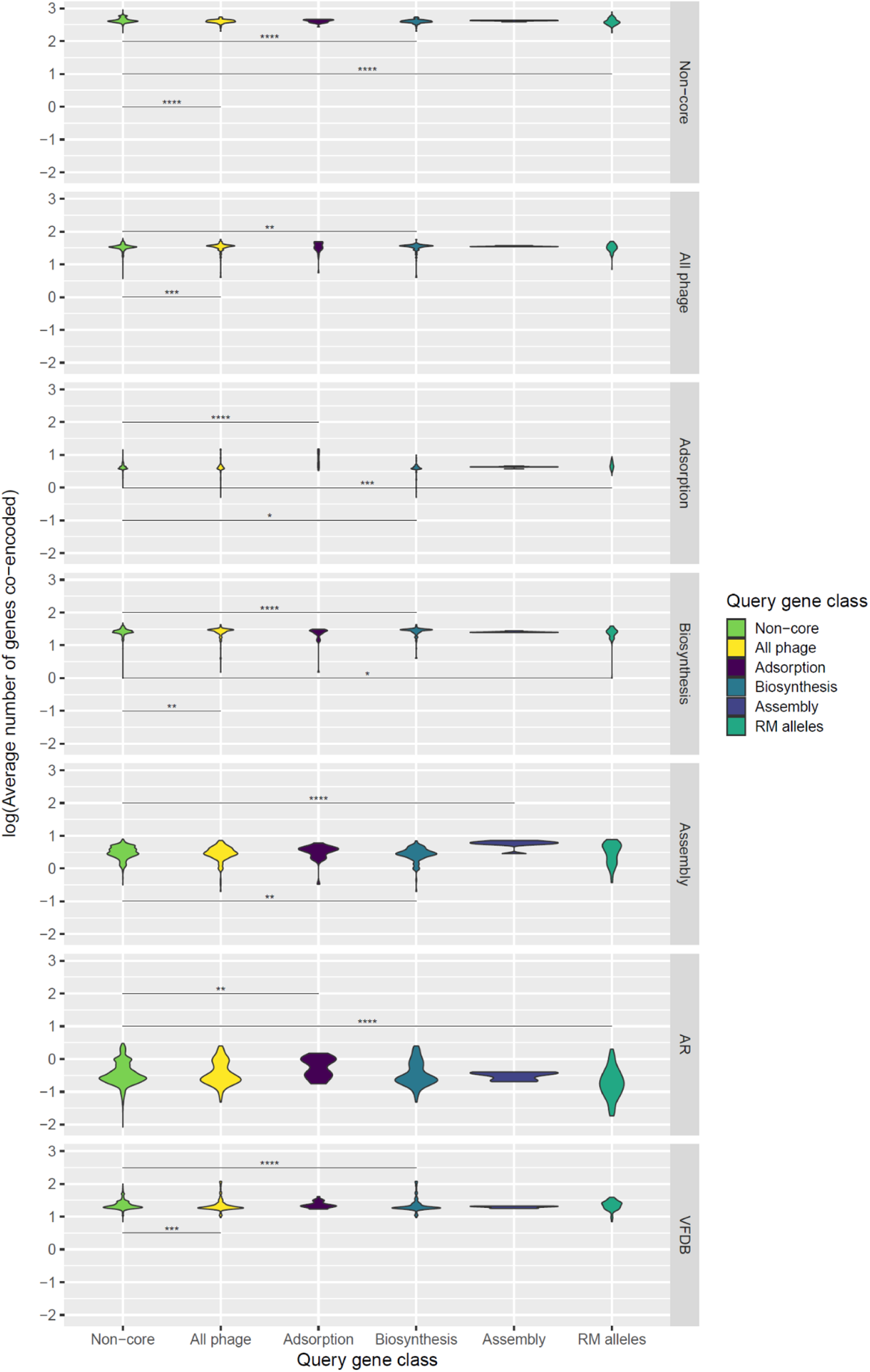
Non-core phage resistance genes are encoded together based on calculating genomic overlaps from a search of 40,000+ *S. aureus* genomes. We used BLAST to search for our non-core (found in less than 80% of genomes) phage resistance genes in the Staphopia database. We then calculated the genomic overlap as the average number of genes in a subject category for strains encoding a query gene. Genomic overlap (y-axis) distributions for different query sets (x-axis) and subject sets (facets) are visualized as violin plots. Group differences relative to all non-core genes were tested for significance with non-parametric Wilcoxon tests (ns, not significant; *, *P* = 0.01 to 0.05; **, *P* = 0.001 to 0.01; ***, *P* = 0.0001 to 0.001; ****, *P* = 0 to 0.0001). AR refers to antibiotic resistance genes while VFDB refers to virulence genes.

### Superinfection immunity correlated with empirically determined phage resistance and accessory genome content

We next asked whether phage resistance gene presence could explain two relevant phenotypes - empirical resistance to infection against 8 phages in 259 diverse *S. aureus* strains determined by laboratory assays (35), and variation in the total number of accessory genes in the set of completely assembled *S. aureus* genomes. While no phage resistance gene categories correlated with measured virulent (p002y and pyo) phage resistance, all phage resistance genes, biosynthesis genes, and superinfection immunity genes (a subset of biosynthesis genes responsible for lysogens repressing superinfecting phages) (36) correlated positively with temperate (p11 and p0040) phage resistance (Figure 6). This result confirmed the hypothesis that prophages confer temperate phage resistance through superinfection immunity. We also saw that total accessory genome content was positively correlated with superinfection immunity genes (R^2^ = 0.26; Figure 7) while all phage resistance genes had only a weak positive correlation (R^2^ = 0.0068). This finding suggested prophages correlated with accessory genome, as strains subject to extensive transduction would potentially also be expected to carry more prophages given the necessary exposure to temperate phage. However, superinfection immunity did not correlate with non-core adsorption, non-superinfection immunity biosynthesis, or assembly gene counts (Supplemental Figure S3), suggesting none of these factors prevented acquisition of superinfection immunity genes through lysogeny. We did note that adsorption correlated slightly with decreased accessory genome content (Figure 7; R^2^ = 0.19).

**Figure 6:**
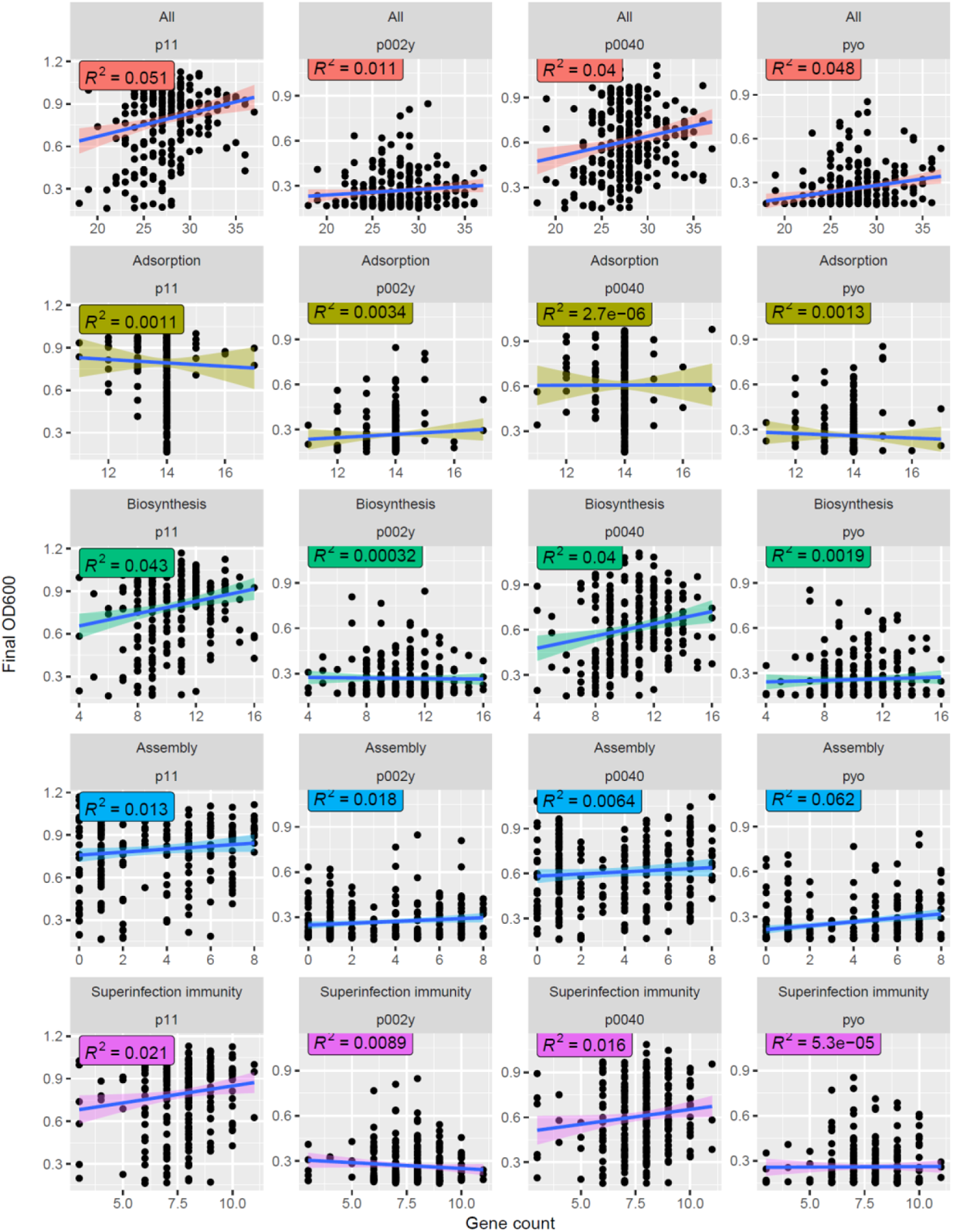
Superinfection immunity but neither adsorption nor assembly genes correlates with empirical temperate phage resistance. We used BLAST to search for our set of non-core phage resistance genes in the set of 263 *S. aureus* genomes from our recent phage host range study. We then plotted the number of matches to all non-core phage resistance, adsorption, biosynthesis, assembly, or superinfection immunity genes (x-axis) against previously measured phage resistance phenotypes (OD_600_ or turbidity after co-culture; y-axis) and calculated correlations (R^2^) between each. We present results for two temperate phages (p11 and p0040) and two virulent phages (p002y and pyo).

**Figure 7:**
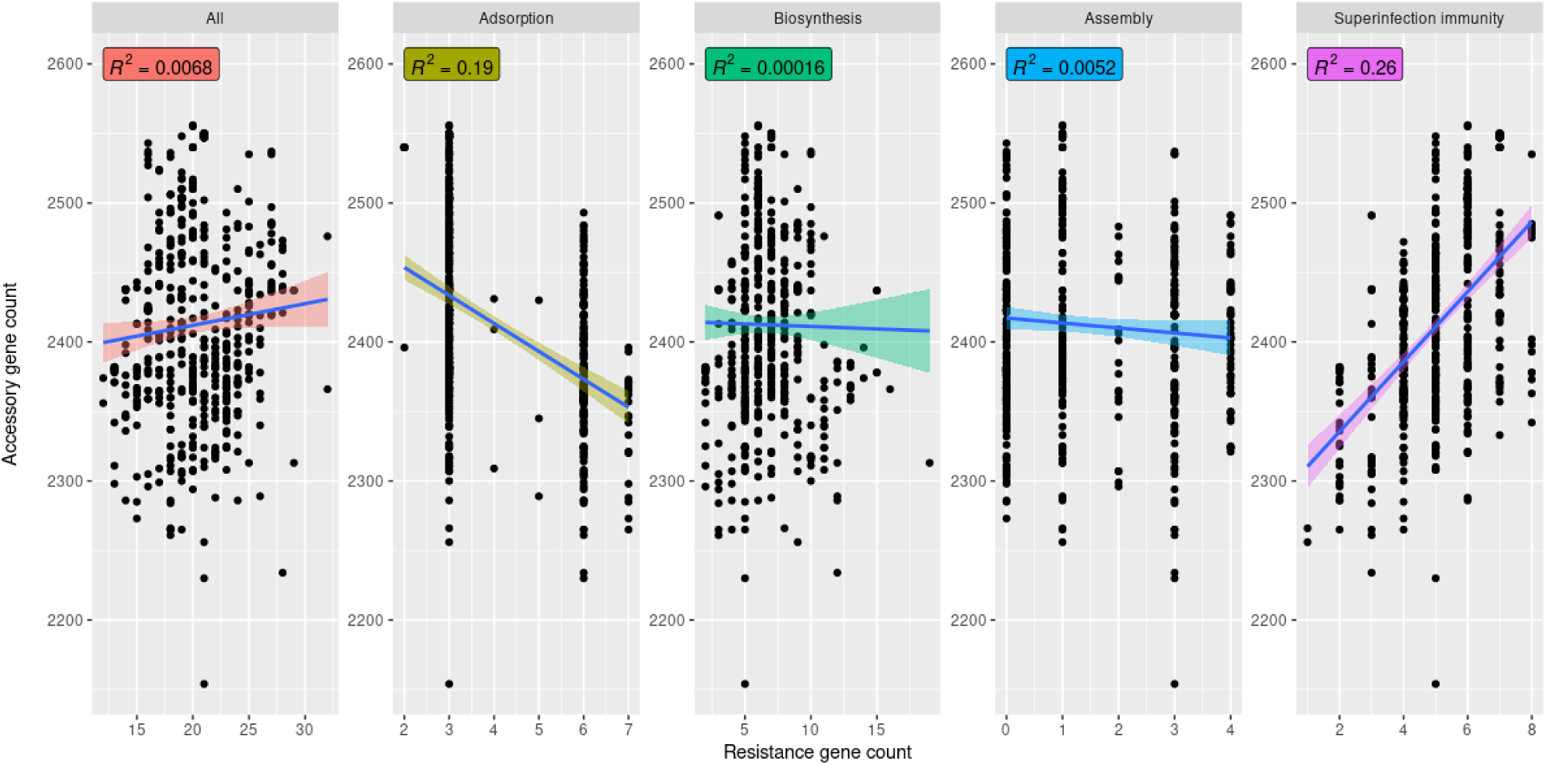
Superinfection immunity but neither adsorption nor assembly genes correlates with accessory genome content. We used BLAST to search for our set of non-core phage resistance genes in the set of 535 complete *S. aureus* genomes in Staphopia. We then plotted the number of matches to all non-core phage resistance, adsorption, biosynthesis, assembly, or superinfection immunity genes (x-axis) against accessory genome content (y-axis) and calculated correlations (R^2^) between each.

To account for quantitation effects of multi-gene systems we compressed matches from gene to system (counting a “system” like restriction-modification to be present if a single system gene was present, such as *hsdR*). We also accounted for the small number of genes that act to reduce phage resistance by multiplying gene presence by a resistance effect factor (1 if gene presence conferred resistance, −1 if gene presence conferred sensitivity). We largely found the same patterns as we did without these corrections (Supplemental Figures S4-S9). Unlike the previous accessory genome analysis, however, we found a positive correlation between biosynthesis gene count, number of systems, or effect and accessory genome content (Supplemental Figures S7-S9; R^2^ = 0.19, 0.097, and 0.19, respectively) when we considered data from the gene presence-absence matrix.

**Figure 9:**
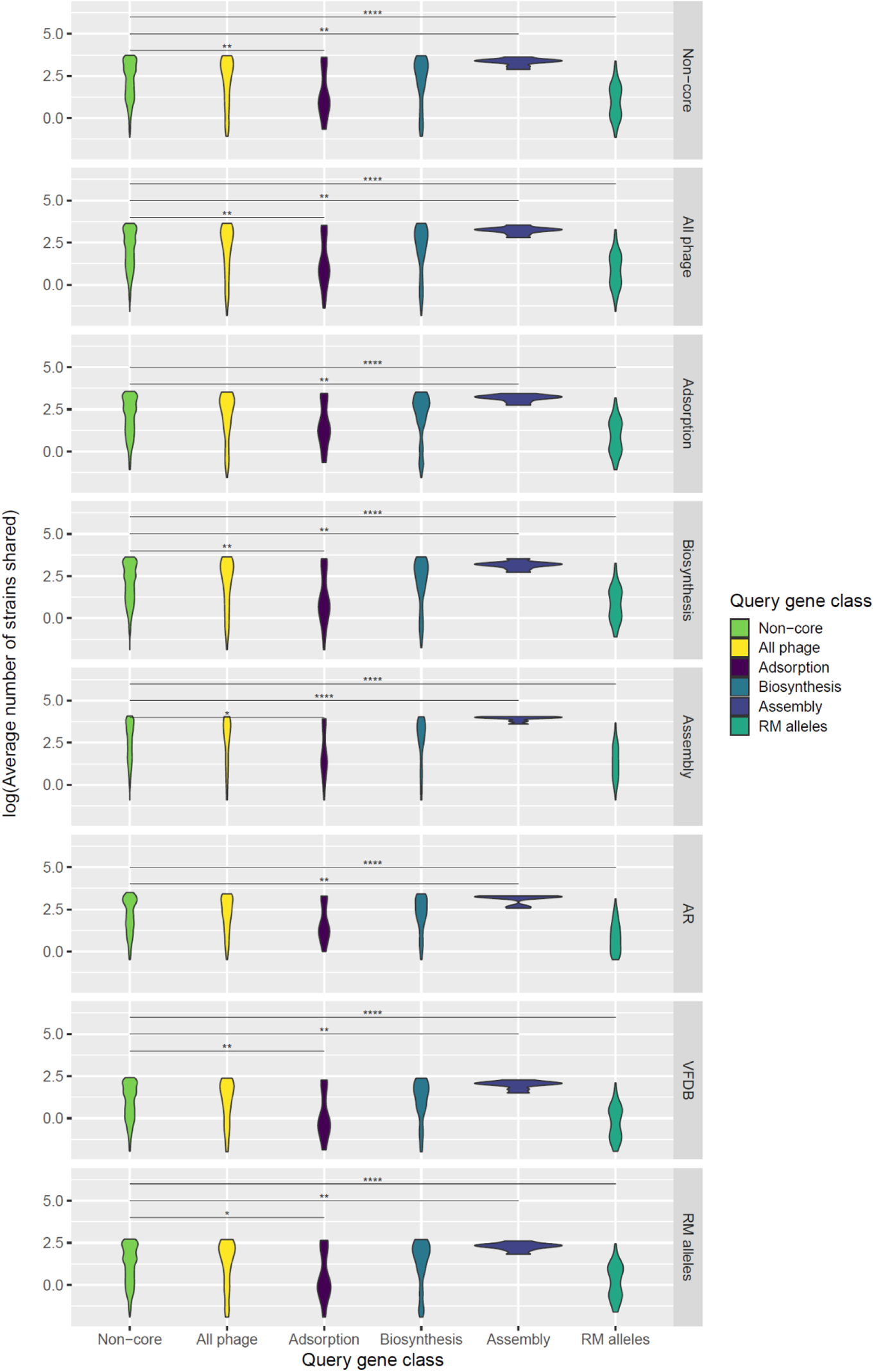
Phylogenetic overlap between classes of non-core genes determined from a search of 40,000+ *S. aureus* genomes. We used BLAST to search for our non-core (found in less than 80% of genomes) phage resistance genes in the Staphopia database. We then calculated the phylogenetic overlap as the average number of encoding strains shared between a query gene and those of a subject category. Phylogenetic overlap (y-axis) distributions for different query sets (x-axis) and subject sets (facets) are visualized as violin plots. Group differences relative to all non-core genes were tested for significance with non-parametric Wilcoxon tests (ns, not significant; *, *P* = 0.01 to 0.05; **, *P* = 0.001 to 0.01; ***, *P* = 0.0001 to 0.001; ****, *P* = 0 to 0.0001). AR refers to antibiotic resistance genes while VFDB refers to virulence genes.

### Relationship between non-core phage resistance genes and horizontal gene transfer, antibiotic resistance, and virulence

Phages are a major conduit for gene acquisition in *S. aureus* and therefore it is possible that strains with greater phage resistance may have reduced uptake of horizontally-transferred virulence and antibiotic resistance genes. We asked whether phage resistance acted as a barrier to antibiotic resistance and virulence gene acquisition by 1) correlating counts of phage resistance genes with non-core antibiotic resistance and virulence gene counts 2) calculating phylogenetic overlap between phage resistance genes of each category and non-core genes or subsets of antibiotic resistance or virulence genes, and 3) calculating genomic overlap between the same sets as 2). We found that total non-core phage resistance and biosynthesis genes positively correlated (R^2^ = 0.072, p = 2.9e-10; R^2^ = 0.099, p = 8.4e-14) with the non-core antibiotic resistance genes, but all categories positively correlated (R^2^ = 0.18, p < 2.2e-16; R^2^ = 0.43, p < 2.2e-16; R^2^ = 0.047, p = 3.6e-07; R^2^ = 0.15, p < 2.2e-16; R^2^ = 0.023, p = 4.4e-4 for all, adsorption, biosynthesis, assembly, and superinfection immunity genes, respectively) with non-core virulence genes (Figure 8). In terms of phylogenetic overlap, assembly genes were co-encoded with antibiotic resistance and virulence genes in more strains than non-core genes on average, while virulence genes were co-encoded with adsorption genes in far fewer strains than non-core genes on average (p<0.05, Wilcoxon test, Figure 9). In all other cases (e.g., non-core genes, all phage resistance genes as subjects), query adsorption genes had less and query assembly genes had more overlap with subject genes than query non-core genes in general (p<0.05, Wilcoxon test, Figure 9). Regarding genomic overlap, more antibiotic resistance genes were encoded for adsorption gene-encoding strains than non-core gene-encoding strains, while more virulence genes were encoded for all phage resistance and biosynthesis gene-encoding strains than non-core gene-encoding strains (p<0.05, Wilcoxon test, Figure 5). These phylogenetic and genomic overlap results thus indicate non-core antibiotic resistance genes tended to be encoded together with adsorption genes, while non-core virulence genes tended to be encoded together with biosynthesis genes.

**Figure 8:**
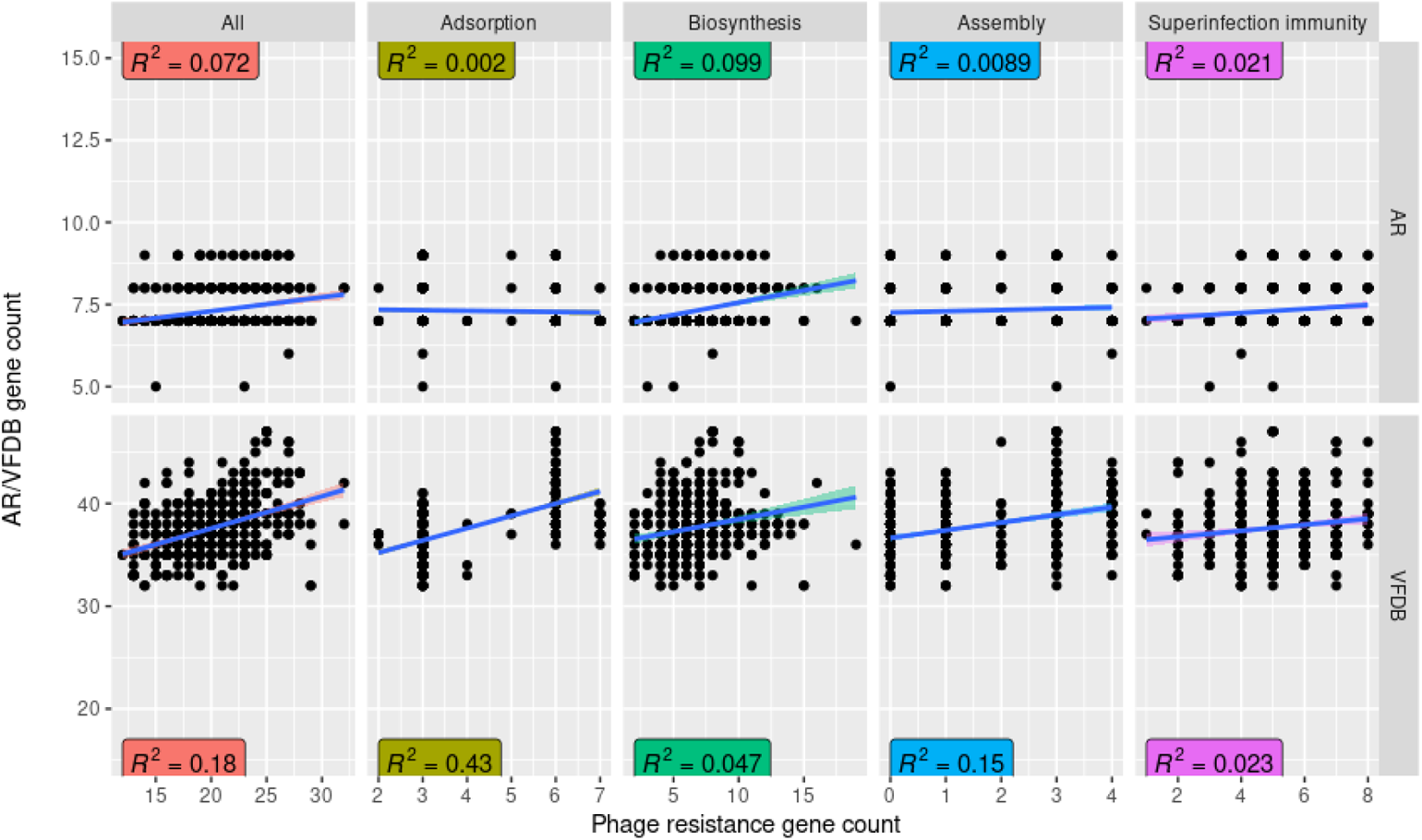
Correlations between non-core antibiotic resistance or virulence genes and non-core phage resistance genes. AR refers to antibiotic resistance genes while VFDB refers to virulence genes. We used BLAST to search for our set of non-core phage resistance, antibiotic resistance, and virulence genes in the set of 535 complete *S. aureus* genomes in Staphopia. We then plotted the number of matches to all non-core phage resistance, adsorption, biosynthesis, assembly, or superinfection immunity genes (x-axis) against matches (y-axis) to non-core antibiotic resistance and virulence genes (facets) and calculated correlations (R^2^) between each.

### Consequences of type I restriction-modification specificity gene (hsdS) allelism on horizontal gene transfer, antibiotic resistance, and virulence

Type I restriction systems are well known as a significant barrier to HGT in *S. aureus* (37, 38). Although core genes, the barrier is dependent on DNA binding specificity of their hsdS DNA binding domain allele. We analyzed type I restriction-modification specificity gene (*hsdS*) alleles for their relationships with HGT, given their correlation with clonal complex and role in restricting transduction between clonal complexes based on restriction specificity (37). We calculated phylogenetic and genomic overlaps between detected NRD *hsdS* alleles (81 total) and all non-core genes or respective antibiotic resistance and virulence gene subsets. In all cases, *hsdS* allele phylogenetic overlap was significantly less (p<0.05, Wilcoxon test) than that of non-core genes generally (Figure 9). However, only for subject non-core, adsorption, biosynthesis, and antibiotic resistance genes were *hsdS* allele genomic overlap significantly different (p<0.05, Wilcoxon test) than that of non-core genes generally (Figure 5). Taken together, these results indicated, as expected, a role for restriction specificity in affecting horizontal gene transfer, especially given that the differences were most pronounced when viewed on the phylogenetic (strains shared between *hsdS* alleles and accessory genes of interest) rather than genomic (number of genes encoded in strains with an *hsdS* allele) level. This is consistent with the hypothesis that phylogeny (CC)-associated *hsdS* alleles restrict horizontal gene transfer.

## Discussion

In this work, we curated genes potentially influencing phage resistance in *S. aureus*, and mapped them against the thousands of genomes in the Staphopia database. We then used these results to ask questions about the relation of genes to empirically determined phage resistance and horizontally transferred antibiotic and virulence genes. The genes were assigned to three stages of phage infection (adsorption, biosynthesis, and assembly) at which *S. aureus* is known to have developed resistance to phages. Adsorption genes examined include wall teichoic acid (WTA) and capsule biosynthesis genes. Biosynthesis genes examined include characterized restriction-modification, abortive infection, and CRISPR systems, as well as newly discovered systems such as cyclic oligonucleotide-based anti-phage signaling systems (CBASS), defense island systems associated with restriction modification (DISARM), retrons, and Lamassu systems, amongst others. This study is the first to find recently discovered Septu, AVAST, and retron phage defense systems (32) in *S. aureus*. Assembly genes examined include all three main mechanisms known in SaPIs - capsid remodeling, packaging interference, and helper phage late gene repression.

We noted four major findings from these analyses. First, we found that core phage resistance genes do not significantly differ in diversity, functionality, and selection from other core genes, refuting a possible arms race hypothesis in the evolution of host receptors and phage. This suggested a different evolutionary dynamic to that seen in *E. coli*, where phages specific to strains in species often exist in an arms race dynamic with the outer-surface of the bacterium, in which phage and host coevolve to outcompete each other (39, 40). Host receptors and phage receptor-binding proteins undergo diversifying selection during this coevolution. We do not see this pattern in *S. aureus*, however, given that core phage resistance genes have similar high functionality (low delta-bit score), low diversity, and negative selection to core genes. We attribute this result to fitness costs of losing capsule and wall teichoic acid (WTA) that these core genes are responsible for synthesizing. Wall teichoic acid is critical for cell division (41–43), methicillin resistance (44), nasal colonization (45), and antimicrobial peptide resistance (46), amongst other roles. Alternatively, our strain set may not capture transient strains resistant through core gene mutation (especially unstable mutations as in phase variation) (47), or non-core genes may instead be undergoing arms race dynamics through frequent gain or loss (48).

A second finding was that non-core phage resistance genes had less phylogenetic signal (measured using consistency index) than other non-core genes. Frequent loss or HGT of these genes or a preponderance of intermediate frequency phage resistance genes would explain this finding. Of the classes of resistance genes, those in the assembly category had the lowest consistency. This may be expected as genes found on prophages (e.g., superinfection immunity determinants) are in this category and frequent HGT of prophages through SaPIs and transduction may lead to inconsistencies with the core genome phylogeny.

Third, we found that non-core phage resistance genes within the same categories are encoded together (e.g., adsorption) by measuring genomic overlap. Thus, if a strain encoded a non-core adsorption gene, for example, it was more likely to encode a gene of the same category than a strain lacking this gene. We note that we did not ask explicitly about genes being encoded together as operons, though recent studies have discovered new phage defense systems based on proximity (but not operonic linkage) to existing systems (29). As was previously noted (29, 32), we also found that non-core recently discovered phage resistance systems were sometimes located near restriction-modification genes. This is likely due to as yet functional interactions between different genes (i.e., one gene depends on another for carrying out a common function) preventing loss of either and transfer of such genes together on common genetic elements (e.g., SaPIs carrying assembly interference genes). In the future, we will examine phage defense islands that contain multiple defense genes and systems in the same genomic region to further address this clustering.

The fourth major finding was that superinfection immunity amongst non-core phage resistance genes was the sole class to correlate with empirical (temperate) phage resistance, accessory genome content, and accessory antibiotic resistance/virulence gene content. We note that superinfection immunity by indicating prophages could correlate with lateral transduction in the pac-type phages, which is worth further study population-wide. Having certain prophages (i.e., pac-type) in the genome could enhance transduction of adjacent genomic DNA several orders of magnitude up to 300 kb downstream, according to recent studies (49).

After discovery in this analysis of some species-wide patterns of phage resistance, the future challenge is to connect them to phenotypes and evolutionary processes. Somewhat surprisingly, simple models of phage resistance gene presence did not predict experimentally determined phage resistance nor measures HGT such as virulence and antibiotic resistance accessory gene numbers. There are several possible reasons why the situation may be more complicated than expected. We cannot expect the presence of a phage resistance gene to correlate with resistance to all phages, when some genes may not affect all phage types and phages are known to have defenses against specific barriers (e.g., anti-restriction and anti-CRISPR systems) (50, 51). It is possible that existing studies characterizing phage resistance genes in *S. aureus* do not reflect growth conditions (e.g., rich medium vs. blood or skin) or selection pressures in natural environments where *S. aureus* is present, making laboratory-defined genes poor predictors of phage resistance in the environment in some cases. It may be that cryptic phage resistance loci not yet characterized are important in predicting each phenotype. It is possible that complete resistance to virulent phage infection is extremely rare and instead subtle metabolic or surface protein changes are instead responsible for most variation in host range. On the other hand, we simply may need to adjust our approach to ignore genes with neutral (type 5 or 8 capsule) or negative (WTA) resistance effects or focus on rare, mobile phage defense systems. Recent studies indicate that rapid turnover of phage defense systems carried on mobile genetic elements explains arms race dynamics between phages and hosts, with the gradient of systems carried creating a gradient of phage sensitivities, at least in the marine *Vibrio* strains examined (48). We may also want to further examine genes for which genetic diversity has effects on phage resistance, such as type I restriction specificity. We also examined phylogenetic overlap between *hsdS* alleles and various non-core genes of interest, for example, finding significantly lower overlap than for non-core genes in general, and a bimodal distribution with a subset of alleles having lower and higher overlap. These findings support the long-held belief that *hsdS* diversity is a major factor shaping horizontal gene transfer patterns in the species through restriction specificity.

Future efforts from the bioinformatic survey should most likely focus on mobile defense systems and prophage diversity. Combinations of these defense systems may impact host range and horizontal gene transfer phenotypes more strongly than most gene classes considered here. The spectrum from cryptic to complete prophages as well as the types should be classified considering their roles in transduction, resistance, and large-scale levels of horizontal gene transfer that this work brings further to light. We also must continue to conduct laboratory phage resistance studies in the species that focus on what is likely to occur in the natural population. Such work would include further boosting the phage host range GWAS with a large number of diverse strains, to classify as many examples of phage resistance in the species as possible.

Resistance evolution studies should also be done in physiologically relevant conditions (e.g., consequences of within-host evolution, biofilm development, and phage challenge at MOIs common in human *S. aureus* niches). All of this future work, together with what has been described here, will enhance our understanding of how phages shape *S. aureus* evolution.

## Materials and Methods

### Curating list of *S. aureus* phage resistance genes

The list of genes known to influence phage resistance was collated from extensive literature review. We previously searched the literature exhaustively for host genes identified through laboratory evolution or molecular genetics to act at three stages of phage infection (adsorption, biosynthesis, and assembly) (4). Genes selected were thus either 1) reported to have causative effects on phage resistance in a *S. aureus* strain through molecular genetic studies, 2) identified through laboratory selection for phage resistance, 3) reported to have causative effects on phage resistance in other species but had homologs in *S. aureus*, or 4) were reported to have causative effects on phage resistance in other species but did not have reported *S. aureus* homologs (to avoid redundancy). An example of the final criterion was the inclusion of the *Lactococcus lactis* superinfection exclusion (*sie*) uptake resistance gene (52), which is not reported in *S. aureus* nor contains a known *S. aureus* homolog. Supplementary Table S1 lists these genes, coordinates and accessions of the associated sequences, the resistance class and subclass, and literature supporting its inclusion in the list, nucleotide sequence selection, and resistance designation.

### Determining phage resistance gene conservation in the *S. aureus* species

Nucleotide sequences of the curated phage resistance gene list were matched against 42,949 *S. aureus* genomes (the Staphopia database) (26) using a BLASTN (53, 54) search with default parameters except maximum target sequences of 10,000,000 and maximum high scoring pairs of 1. BLAST output was filtered for unique matches between each gene and each strain. We then counted the number of unique matches per gene to determine the number of strains containing each gene. Gene conservation in the species was then compared between phage resistance gene categories (adsorption, biosynthesis, and assembly) using violin plots and assessed for statistically significant differences between groups with non-parametric Wilcoxon tests.

### Visualizing genomic contexts of non-core recently discovered phage resistance systems

In addition to evaluating conservation of phage resistance genes, we examined the genomic surroundings of the most recently discovered phage resistance systems in our set (mainly retrons and cyclic oligonucleotide-based antiphage signaling systems - CBASS) (32, 55). We first constructed the pangenome of the Staphopia completed genome set (535 genomes) with PIRATE (56). We then matched phage resistance gene sequences (Supplemental Table S1) from the two previously cited papers (32, 55) against non-core (present in 80% or less of the genomes) PIRATE pangenome clusters and identified up to the first five genomes containing this non-core gene subset (seven genes total). Five pseudogene or CDS sequences up or downstream of the non-core phage resistance gene of interest were selected from the respective genome GFF file for each genome and each non-core phage resistance gene of interest. Non-core genes and their genomic contexts obtained as described were then visualized using the R package gggenes (57).

### Determining core phage resistance gene diversity, functionality, and selection

The pangenome of the Staphopia non-redundant (NRD) set (380 strains representing each sequence type) (26) was constructed using PIRATE (56) run with default parameters. Core genes were those unique PIRATE gene clusters only present in 80% of NRD genomes (304) or more. We focused gene diversity, functionality, and selection studies on these core phage resistance genes, comparing these sets against corresponding total core genes (excluding core phage resistance). We evaluated gene diversity both through the number of alleles per corresponding gene in the pangenome and the translated nucleotide sequence amino acid diversity (π) calculated from the corresponding gene’s pangenome nucleotide alignment using modified scripts originally written by John Lees (58). We evaluated functionality using delta-bit score (33) and measured selection by calculating dN/dS for each gene with the package Hypothesis Testing using Phylogenies (HyPhy) (59). PIRATE output provided the number of alleles at the maximum cutoff (98%) for each gene, so it was directly parsed to get allele counts for each core gene. Amino acid diversity (π) was calculated from translated PIRATE gene cluster nucleotide alignments. dN/dS was calculated using individual gene phylogenetic trees inferred from nucleotide alignments with IQ-TREE (60, 61). dN/dS was calculated using single-likelihood ancestor counting (SLAC), which uses maximum likelihood (ML) and counting approaches to infer dN and dS per site given a codon alignment and corresponding phylogeny (59, 62, 63). SLAC first predicts the most likely ancestral sequence at each node of the phylogeny with ML and then counts nonsynonymous and synonymous changes per site in a manner similar to the Suzuki-Gojobori method (63, 64). For delta-bit score analysis, for each tested gene, we calculated average delta-bit scores for all strains encoding protein sequences that matched a corresponding HMM. Proteins were matched to Pfam family HMMs using HMMER (65).

### Evaluating phylogenetic associations with non-core phage resistance genes

We evaluated phylogenetic associations with phage resistance genes by 1) determining homoplasy for each non-core phage resistance gene and 2) correlating clonal complex (CC) with non-core phage resistance gene count. Non-core genes were defined as those present in less than 80% of the genomes and filtered for redundancy (unique PIRATE gene cluster matches to query phage resistance genes were selected for further analysis). Homoplasy measurement through consistency index (CI) calculation was conducted with HomoplasyFinder (66) given gene presence/absence input and the NRD set phylogenetic tree. We constructed the NRD set maximum-likelihood phylogenetic tree with IQ-TREE using the gubbins (67)-recombination corrected PIRATE core genome alignment. Consistency index, or the consistency between a character amongst strains and that expected on the tree, was calculated from (number of possible character - 1)/(number of changes necessary to explain the character pattern on the tree) or 1/(number of necessary changes) because only two outcomes were possible for each gene (presence or absence). To assess whether consistency indices were statistically significant, we calculated average CI values and their standard deviations for the original data plus 999 permutations of the gene presence/absence data on the tree. We transformed CI to the number of necessary changes (1/CI) for better data visualization and further comparisons. We compared the number of changes for non-extended-core genes to non-extended-core phage resistance genes of each category with violin plots and assessed significance with non-parametric Wilcoxon tests. We also plotted the number of necessary changes to explain the character pattern against the number of strains encoding the gene to determine a relationship between these factors and to compare the relationships for the actual and permuted data. In addition, we also compared non-core phage resistance gene count by clonal complex in the complete genome set (535 genomes). We compared counts for each phage resistance category along with accessory genome content using a boxplot and assessed statistical significance of overall differences with an analysis of variance (ANOVA) statistical test.

### Non-extended-core resistance gene correlation analyses with empirical phage resistance and accessory genome content

We examined the relationship between non-extended-core phage resistance genes and 1) experimentally measured phage resistance phenotypes, 2) accessory genome content, and 3) non-extended-core antibiotic resistance or virulence gene content. We used genomes of previously resistance-phenotyped strains from our *S. aureus* genome-wide host range study (35) and genomes of all completely assembled *S. aureus* genomes (26) to address the first and second objectives, respectively (the third we addressed with both sets). Antibiotic resistance genes searched were previously identified in *S. aureus* genomes (68), whereas virulence genes searched were the Virulence Factor Database (VFDB) set (69). As for the NRD set, non-extended-core phage resistance, antibiotic resistance, and virulence genes were defined as those unique PIRATE gene cluster matches present in 80% of the respective genome set (complete or GWAS). We then used BLAST to match these three sets of non-extended-core genes in the complete and GWAS genome sets. BLAST matches were filtered by query coverage relative to subject, only keeping those with 60% or higher. Matches were further filtered for uniqueness. Filtered numbers of matches were then plotted against empirically measured phage resistance (GWAS set), accessory genome content, or non-core antibiotic resistance or virulence gene matches. Linear regressions were performed on each distribution to assess correlations between these variables.

In addition, we performed two subsequent analyses in which we converted phage resistance gene matches to the number of systems and net phage resistance effects. These analyses relied on the PIRATE gene presence-absence matrices (complete or GWAS) rather than BLAST matches to genes of interest. Number of systems per strain was counted as the number of subclasses (Supplemental Table S1) for which there was a match to at least one gene. Net phage resistance effect on the other hand was determined for each strain by multiplying the phage resistance gene presence-absence matrix by a vector containing the direction (Supplemental Table S1) of the presence of each phage resistance gene (+1 for more resistance and −1 for more sensitivity if a gene is present). Number of systems or net phage resistance effect was then plotted against empirically measured phage resistance (GWAS set), accessory genome content, or non-core antibiotic resistance or virulence gene matches. Linear regressions were performed on each distribution to assess correlations between these variables.

### Calculating non-core phage resistance gene phylogenetic and genomic overlap

In addition to assessing correlations between non-core phage resistance gene counts and accessory genome content on a per-strain level, we also evaluated strain and gene level concordance between non-extended-core phage resistance genes. We did this, as for the correlation analysis, to determine associations among classes of phage resistance genes or between such classes and accessory antibiotic resistance or virulence genes, but with phylogenetic or genomic corrections on a much larger dataset. Unlike the previous analysis, we instead searched phage resistance, antibiotic resistance, and virulence (VFDB) genes against our full Staphopia database with BLAST. BLAST matches were filtered by query coverage relative to subject, only keeping those with 60% or higher. Matches were further filtered for uniqueness and converted to a list of strain-gene pairs. Strain-gene pairs were compared to lists of perfect strain-gene pairs (all strains matching to each gene) to convert the list into a presence absence matrix. Only Staphopia non-extended-core genes (phage resistance genes from the complete list present in less than 80% of Staphopia genomes) were considered for further analysis. Genomic overlap per gene was calculated as the average number of genes in a category (e.g., adsorption) encoded by strains encoding the gene of interest, while phylogenetic overlap per gene was calculated as the total number of genes in a category encoded by all strains encoding the gene of interest divided by the number of genes in that category. Genomic overlap measures how many genes of a certain type are co-encoded with a gene of interest on average, while phylogenetic overlap measures how many strains co-encode a gene and all those of a certain type on average. Genomic and phylogenetic overlap were compared between groups with violin plots and significant differences assessed with non-parametric Wilcoxon tests. For the *hsdS* gene, this analysis was repeated with the 81 alleles detected in the NRD set PIRATE pangenome. Genomic and phylogenetic overlap distributions were compared with violin plots and significant differences assessed using non-parametric Wilcoxon tests.

## Acknowledgements

We thank members of the lab - Michelle Su, Emily Wissel, Robert Petit, and Ashley Alexander - for constructive criticism. Timothy D. Read was supported by United States National Institutes of Health (NIH) R21 AI 138079-02, while Abraham G. Moller was supported by the United States National Science Foundation (NSF) Graduate Research Fellowship Program (GRFP).

---

**Supplemental Figure S1:**
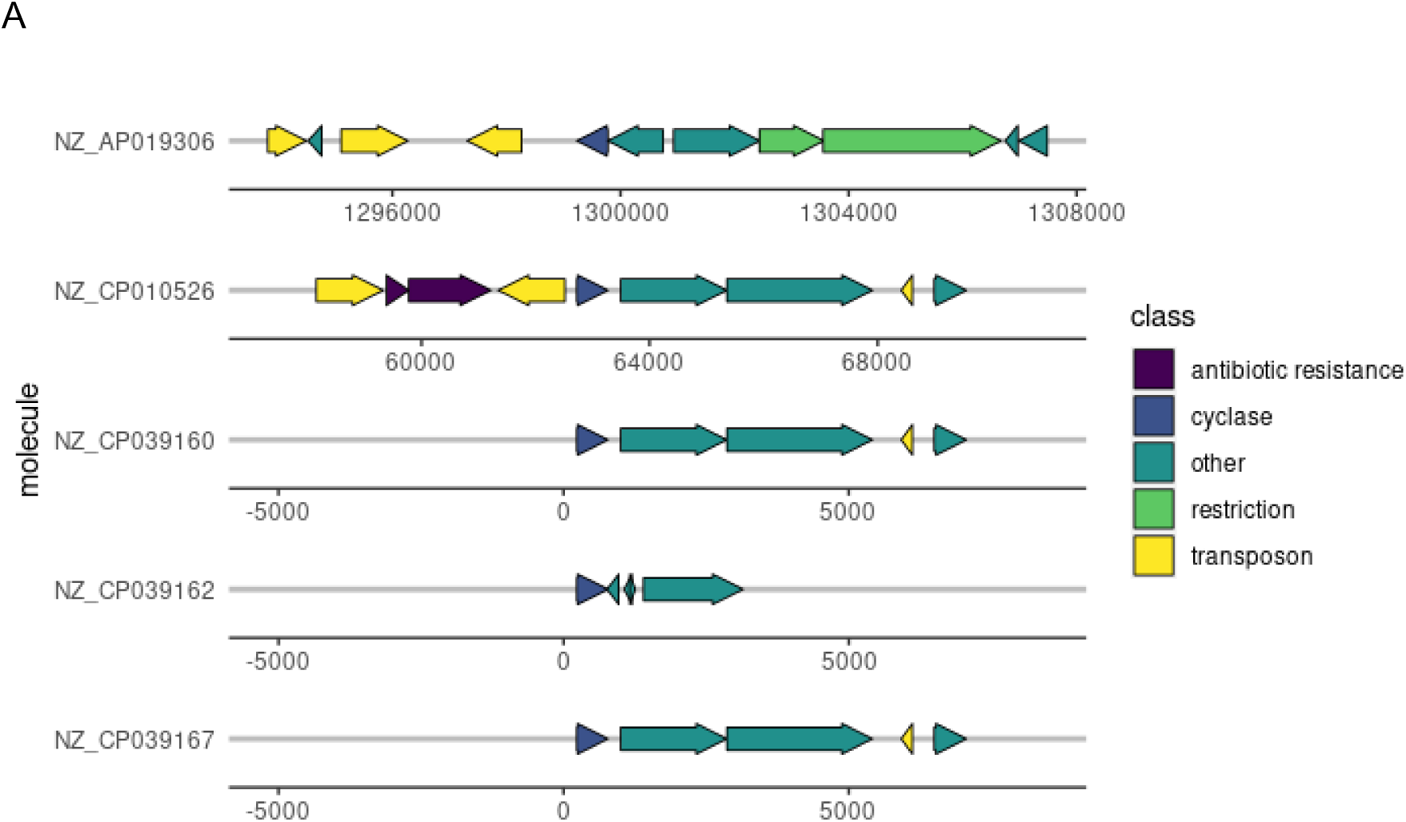
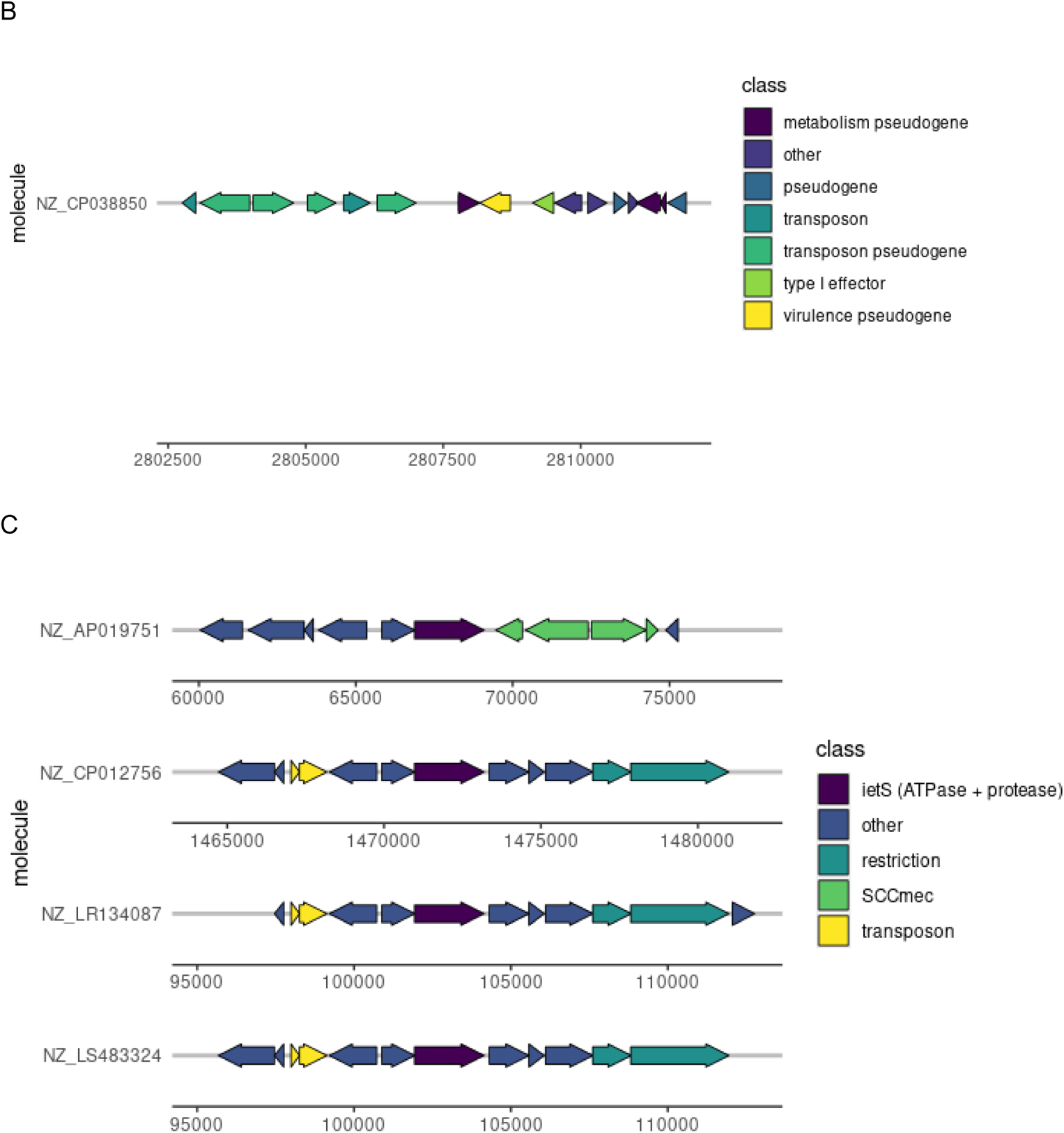
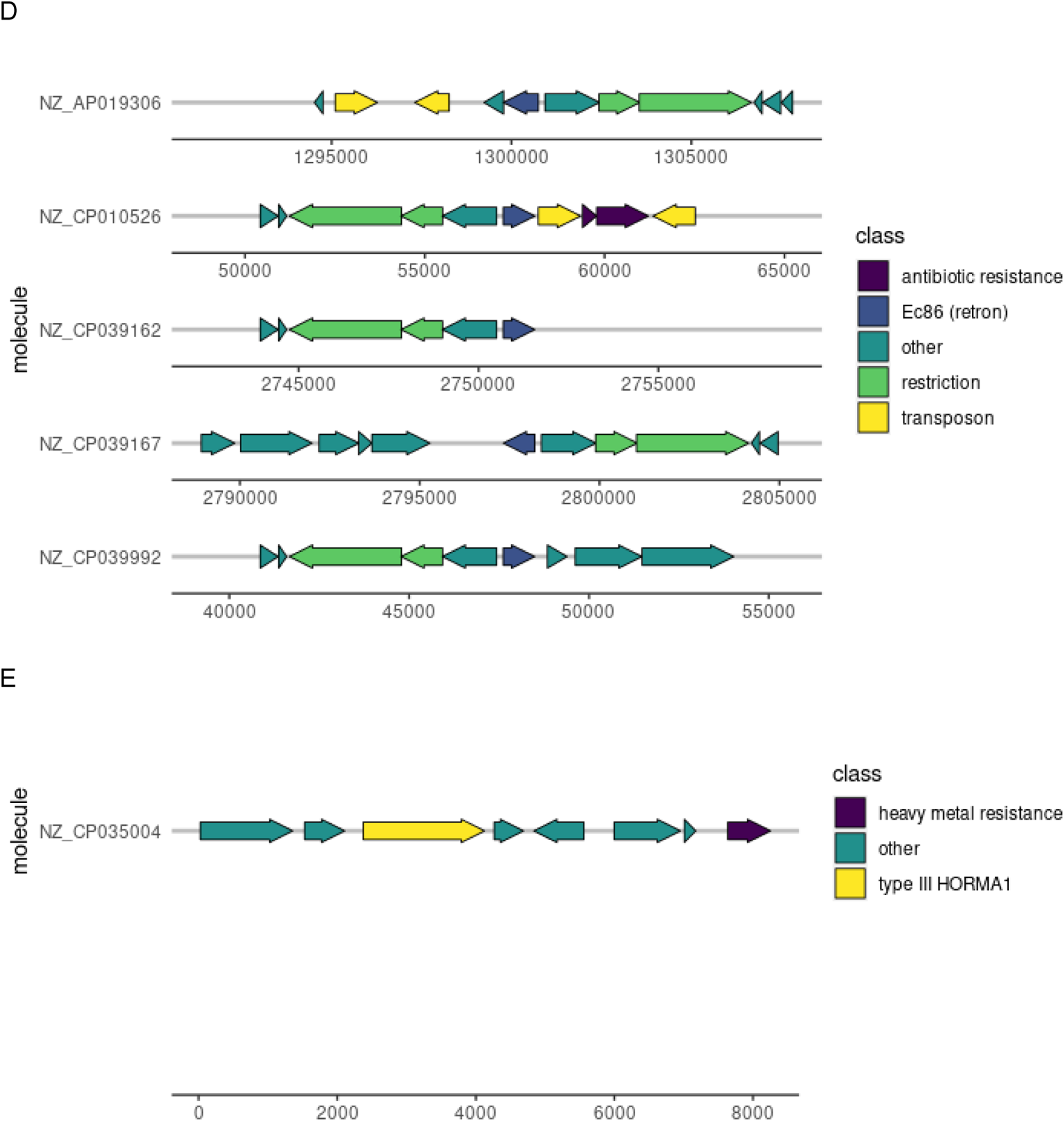
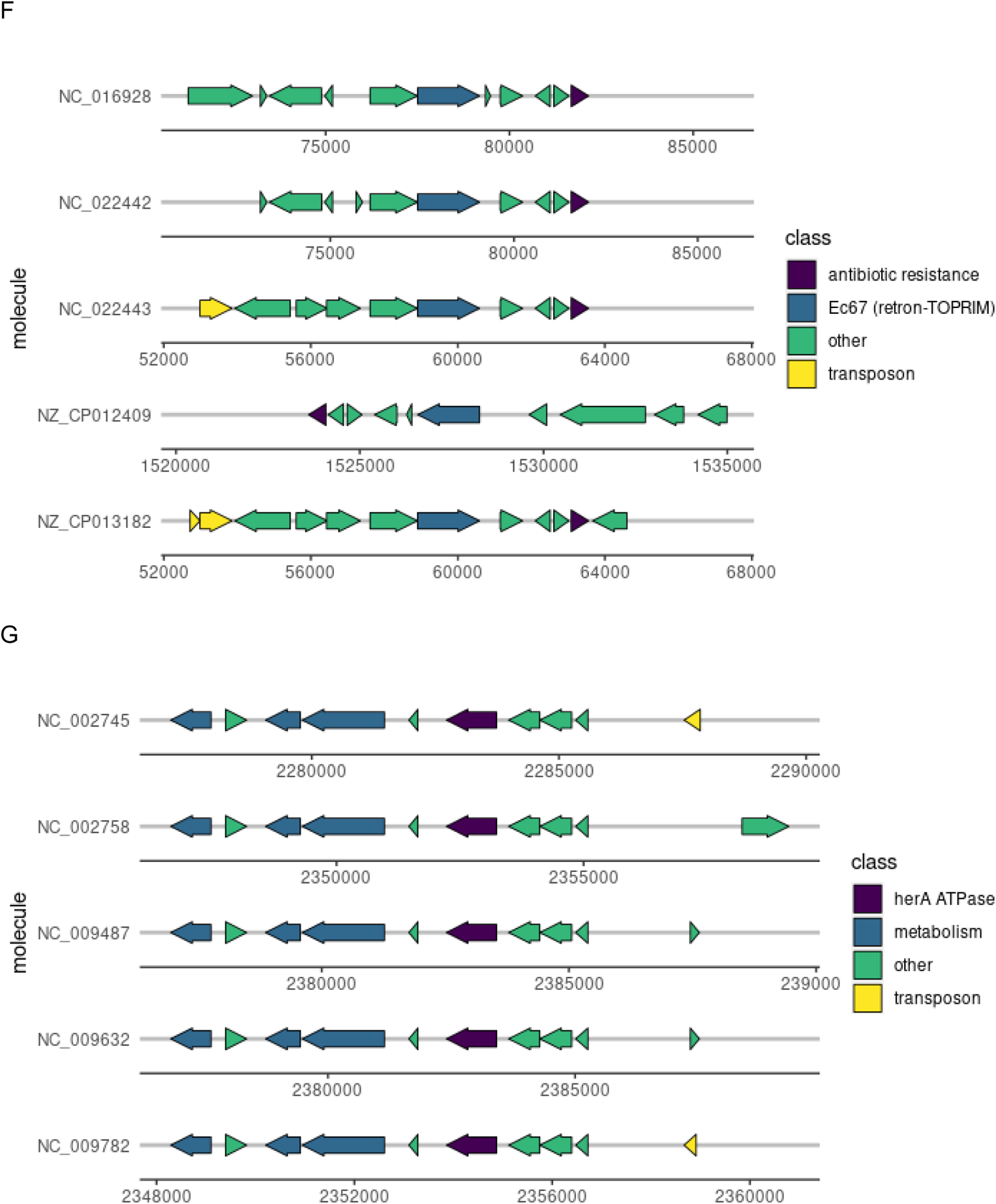
Genomic contexts of non-core phage resistance genes in the Staphopia complete genome set. A-G correspond to the cyclase, type I (cyclase) effector, *ietS*, Ec86 (retron), type III HORMA1, Ec67 (retron-TOPRIM), and *herA* genes, respectively. A maximum of five genomes were selected that contained each non-core phage resistance gene. Five genes upstream and five downstream of the gene of interest were selected for visualization. Genes are colored by common functional category.

**Supplemental Figure S2:**
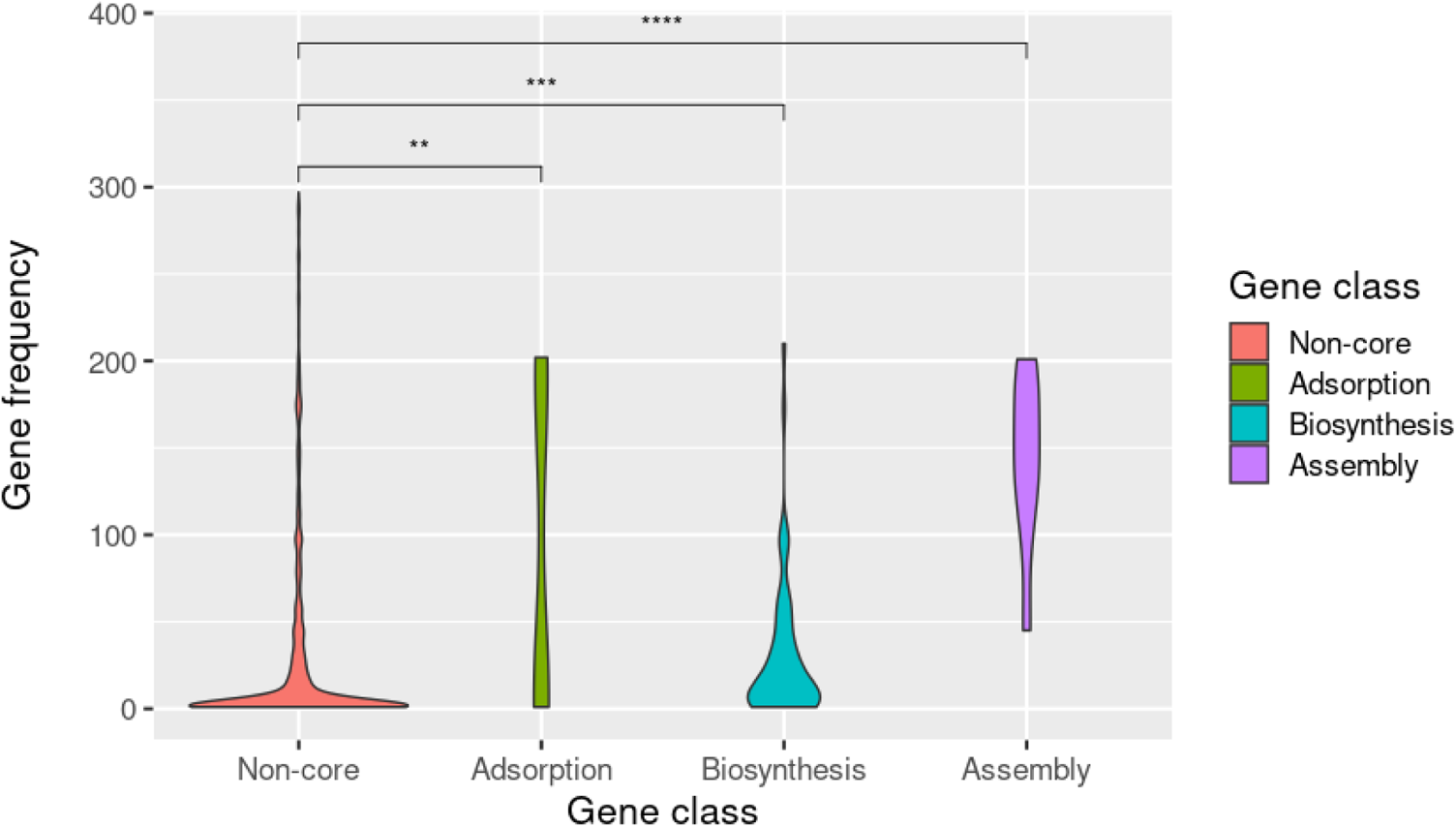
Distributions of gene frequency (number of genomes in set encoding the gene) for each gene category (non-core, adsorption, biosynthesis, and assembly) visualized as violin plots. Group differences were tested for significance using the non-parametric Wilcoxon test (ns, not significant; *, *P* = 0.01 to 0.05; **, *P* = 0.001 to 0.01; ***, *P* = 0.0001 to 0.001; ****, *P* = 0 to 0.0001).

**Supplemental Figure S3:**
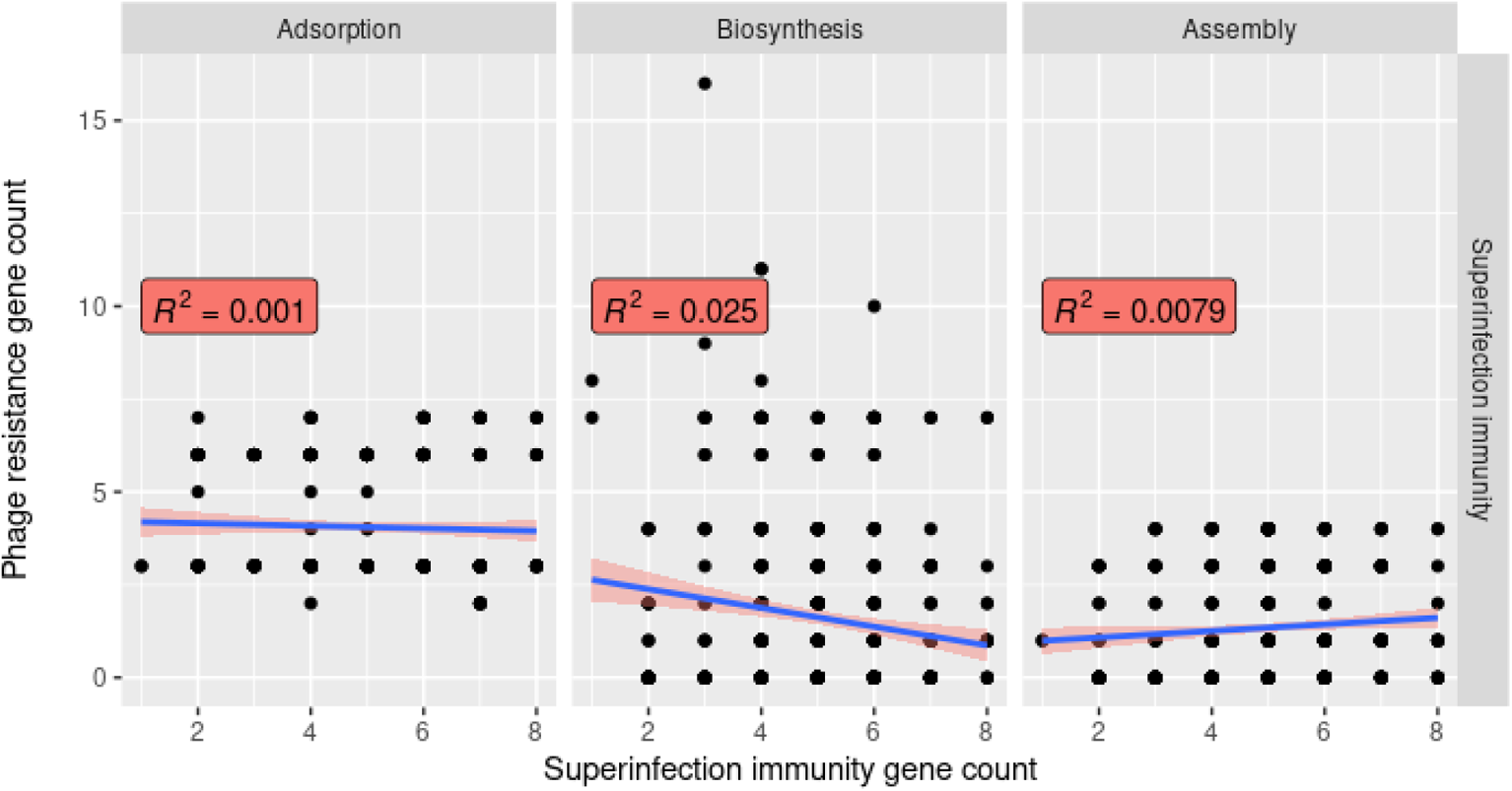
Superinfection immunity does not correlate with non-core adsorption, non-superinfection immunity biosynthesis, or assembly gene counts. We used BLAST to search for our set of non-core phage resistance genes in the set of 535 complete *S. aureus* genomes in Staphopia. We then plotted the number of matches to all non-core adsorption, non-superinfection immunity biosynthesis, or assembly genes (y-axis) against matches to non-core superinfection immunity genes (x-axis) and calculated correlations (R^2^) between each.

**Supplemental Figure S4:**
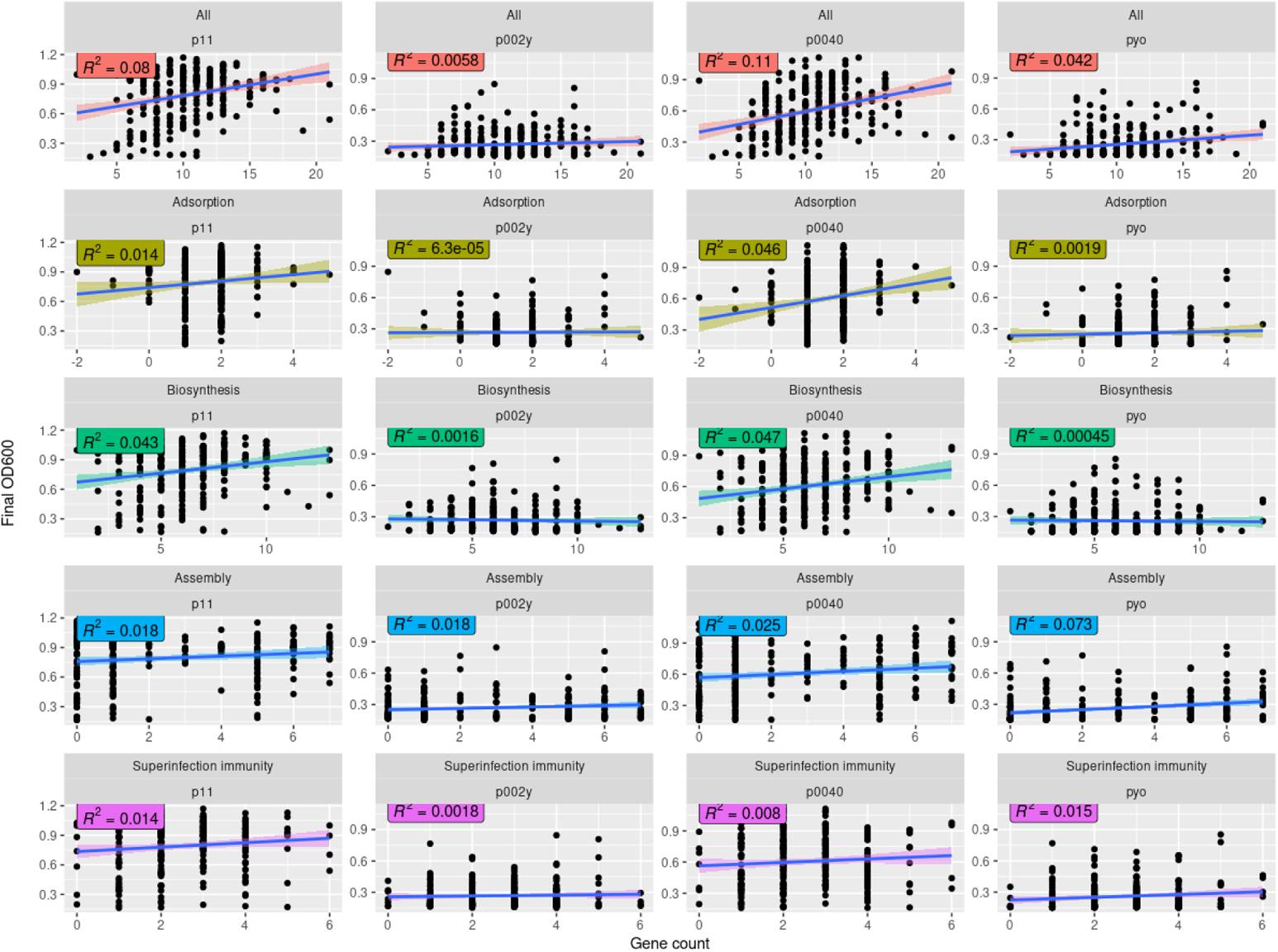
Superinfection immunity but neither adsorption nor assembly genes correlates with empirical temperate phage resistance. We counted non-core phage resistance gene clusters present per genome in our set of 263 genomes with a PIRATE gene presence-absence matrix. We then plotted the number of all non-core phage resistance, adsorption, biosynthesis, assembly, or superinfection immunity genes present (x-axis) against previously measured phage resistance phenotypes (OD_600_ or turbidity after co-culture; y-axis) and calculated correlations (R^2^) between each. We present results for two temperate phages (p11 and p0040) and two virulent phages (p002y and pyo).

**Supplemental Figure S5:**
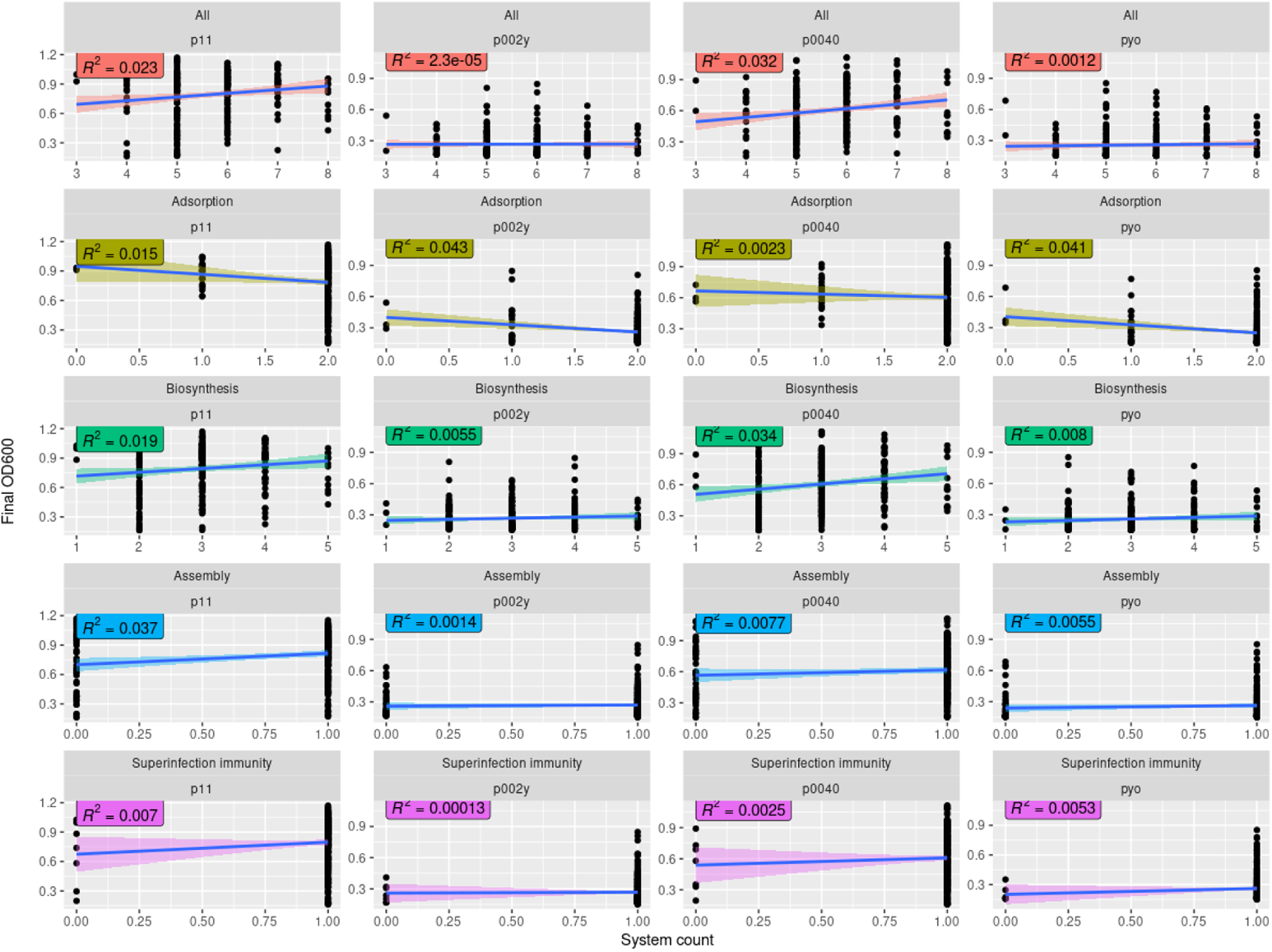
Superinfection immunity but neither adsorption nor assembly genes correlates with empirical temperate phage resistance. We counted non-core phage resistance systems present per genome in our set of 263 genomes with a PIRATE gene presence-absence matrix. We counted a system as present if any individual system gene (e.g., *hsdR* for type I restriction-modification) was present. We then plotted the number of non-core phage resistance, adsorption, biosynthesis, assembly, or superinfection immunity systems present (x-axis) against previously measured phage resistance phenotypes (OD_600_ or turbidity after co-culture; y-axis) and calculated correlations (R^2^) between each. We present results for two temperate phages (p11 and p0040) and two virulent phages (p002y and pyo).

**Supplemental Figure S6:**
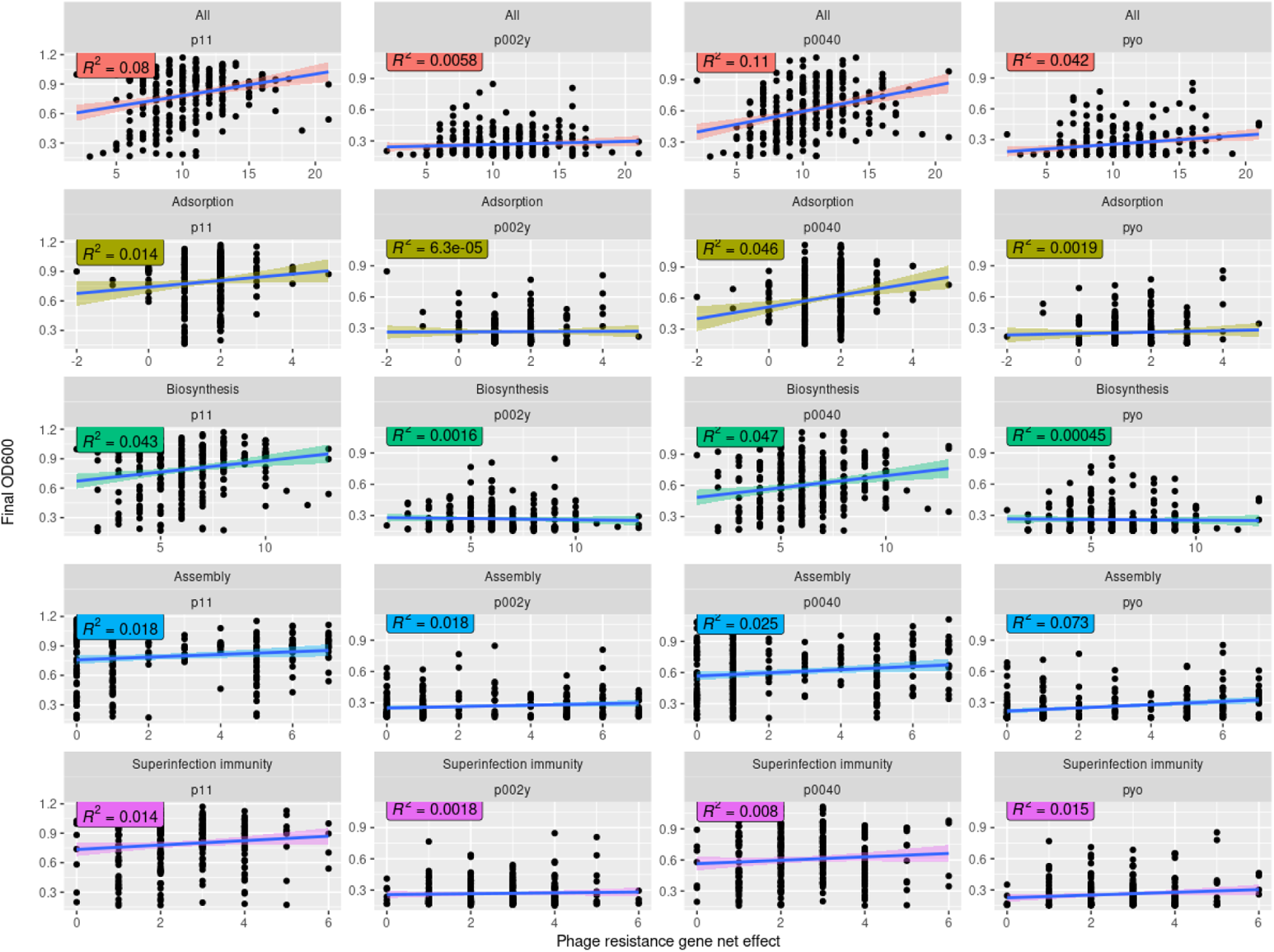
Superinfection immunity but neither adsorption nor assembly genes correlates with empirical temperate phage resistance. We calculated net non-core phage resistance effect per genome in our set of 263 genomes with a PIRATE gene presence-absence matrix. We did so by multiplying the gene presence-absence matrix by a vector indicating effect direction per gene (+1 for gene presence conferring resistance and −1 for gene presence conferring sensitivity) and then calculating the sum of the modified gene presence-absence matrix per strain. We then plotted the non-core phage resistance, adsorption, biosynthesis, assembly, or superinfection immunity net phage resistance effect (x-axis) against previously measured phage resistance phenotypes (OD_600_ or turbidity after co-culture; y-axis) and calculated correlations (R^2^) between each. We present results for two temperate phages (p11 and p0040) and two virulent phages (p002y and pyo).

**Supplemental Figure S7:**
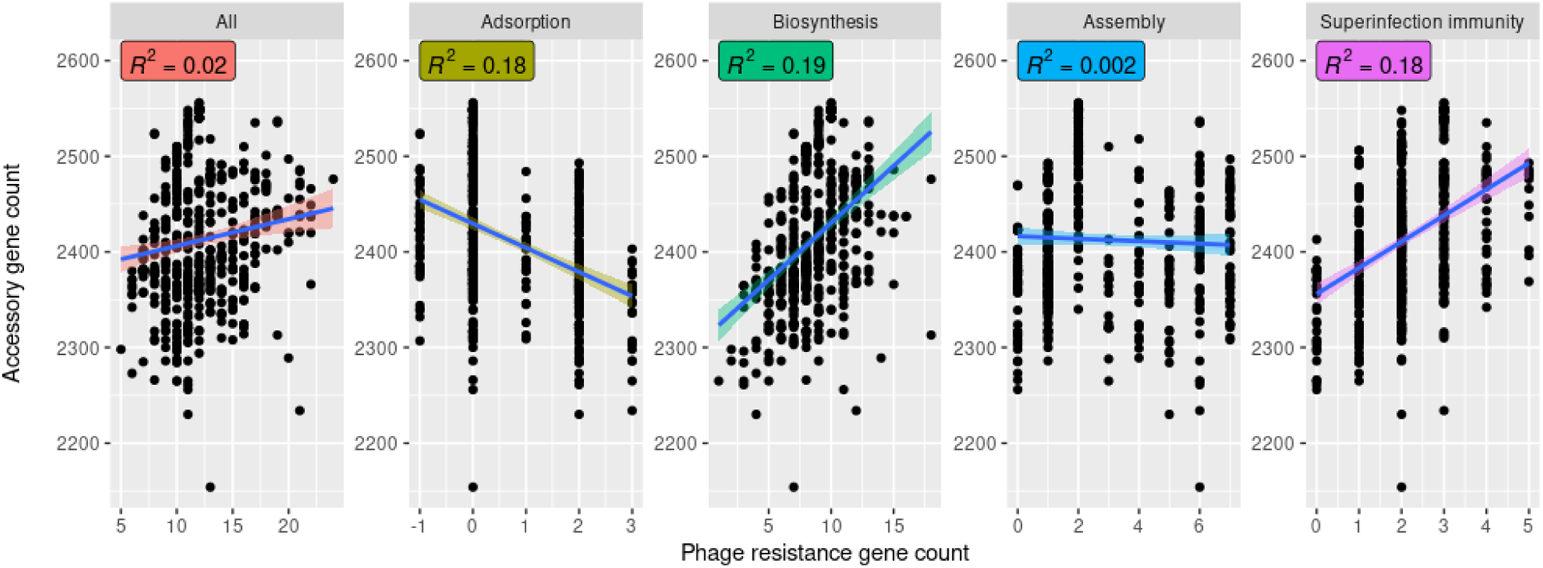
Superinfection immunity but neither adsorption nor assembly genes correlates with accessory genome content. We counted non-core phage resistance gene clusters present per genome in our set of 535 complete *S. aureus* Staphopia genomes with a PIRATE gene presence-absence matrix. We then plotted the number of all non-core phage resistance, adsorption, biosynthesis, assembly, or superinfection immunity genes present (x-axis) against accessory genome content (y-axis) and calculated correlations (R^2^) between each.

**Supplemental Figure S8:**
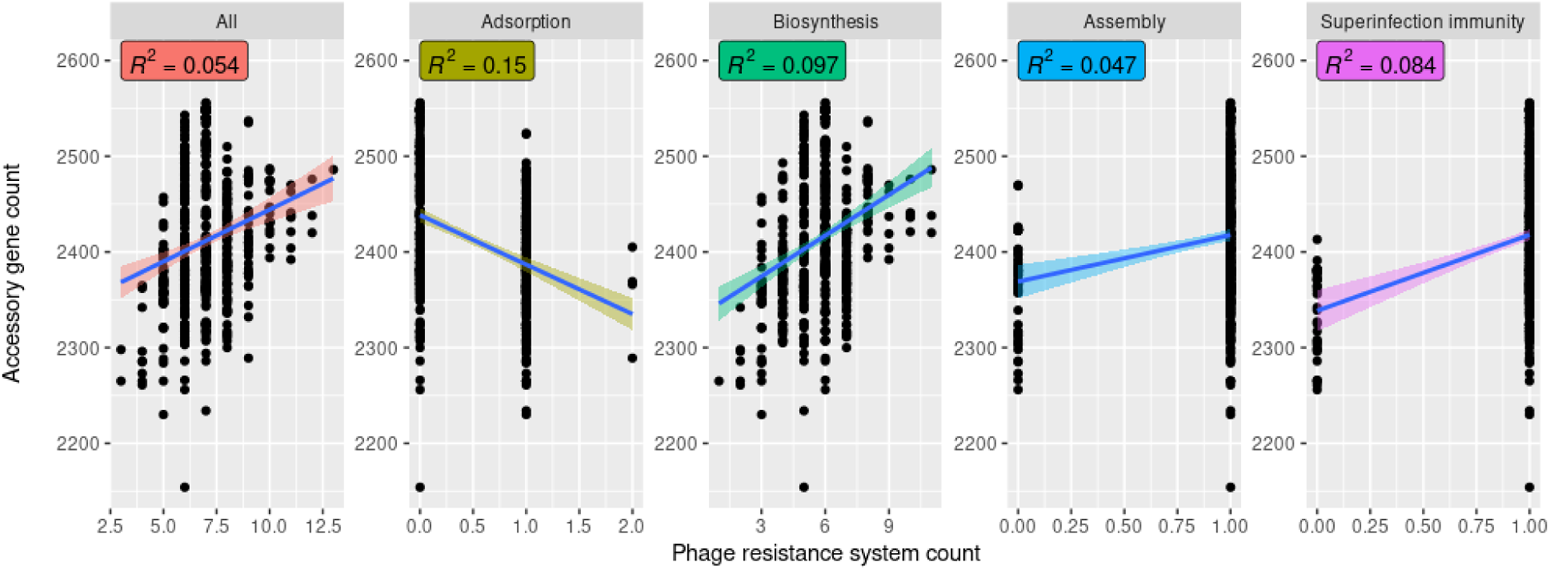
Superinfection immunity but neither adsorption nor assembly genes correlates with accessory genome content. We counted non-core phage resistance systems present per genome in our set of 535 complete *S. aureus* Staphopia genomes with a PIRATE gene presence-absence matrix. We counted a system as present if any individual system gene (e.g., *hsdR* for type I restriction-modification) was present. We then plotted the number of all non-core phage resistance, adsorption, biosynthesis, assembly, or superinfection immunity systems (x-axis) against accessory genome content (y-axis) and calculated correlations (R^2^) between each.

**Supplemental Figure S9:**
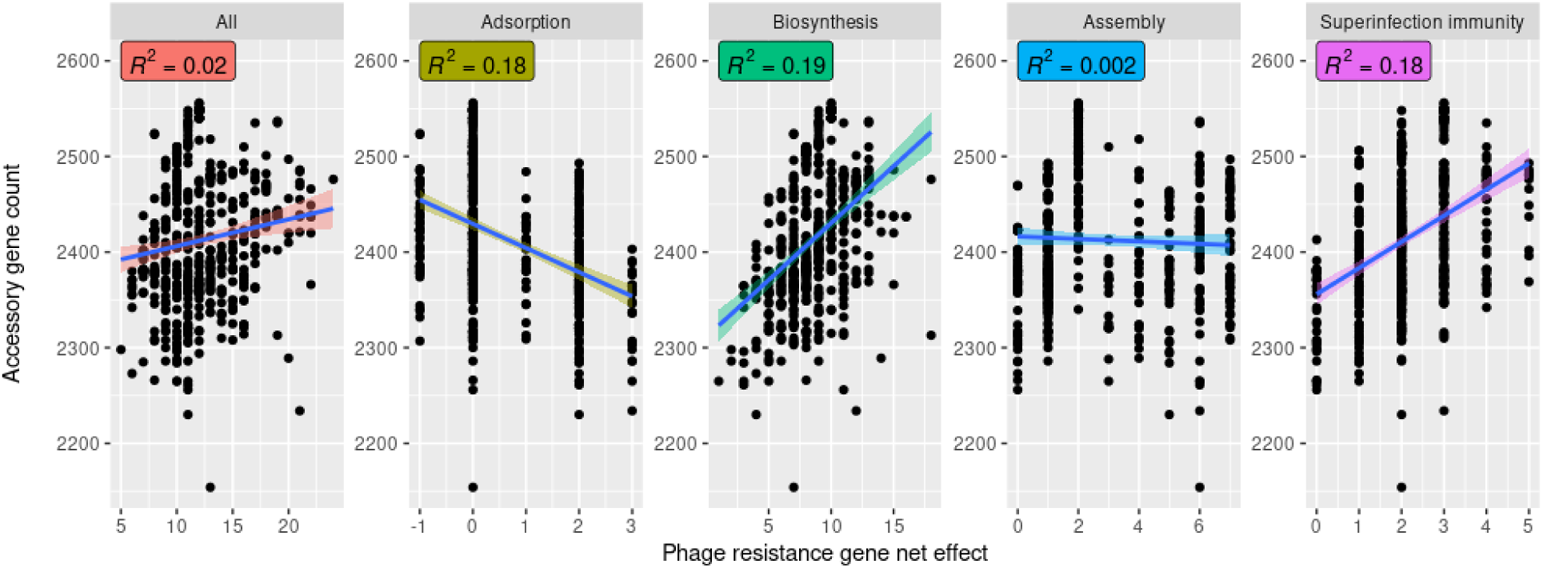
Superinfection immunity but neither adsorption nor assembly genes correlates with accessory genome content. We calculated net non-core phage resistance effect per genome in our set of 535 complete *S. aureus* Staphopia genomes with a PIRATE gene presence-absence matrix. We did so by multiplying the gene presence-absence matrix by a vector indicating effect direction per gene (+1 for gene presence conferring resistance and −1 for gene presence conferring sensitivity) and then calculating the sum of the modified gene presence-absence matrix per strain. We then plotted the number of matches to all non-core phage resistance, adsorption, biosynthesis, assembly, or superinfection immunity genes (x-axis) against accessory genome content (y-axis) and calculated correlations (R^2^) between each.

Supplemental Table S1: List of curated phage resistance genes with strains, genome coordinates, accessions, classes (e.g., adsorption), and subclasses (e.g., receptor), attached as an Excel spreadsheet.

